# The evolutionary history of hepaciviruses

**DOI:** 10.1101/2023.06.30.547218

**Authors:** YQ Li, M Ghafari, AJ Holbrook, I Boonen, N Amor, S Catalano, JP Webster, YY Li, HT Li, V Vergote, P Maes, YL Chong, A Laudisoit, P Baelo, S Ngoy, SG Mbalitini, GC Gembu, P Musaba Akawa, J Goüy de Bellocq, H Leirs, E Verheyen, OG Pybus, A Katzourakis, AN Alagaili, S Gryseels, YC Li, MA Suchard, M Bletsa, P Lemey

## Abstract

In the search for natural reservoirs of hepatitis C virus (HCV), a broad diversity of non-human viruses within the *Hepacivirus* genus has been uncovered. However, the evolutionary dynamics that shaped the diversity and timescale of hepaciviruses evolution remain elusive. To gain further insights into the origins and evolution of this genus, we screened a large dataset of wild mammal samples (*n =* 1,672) from Africa and Asia, and generated 34 full-length hepacivirus genomes. Phylogenetic analysis of these data together with publicly available genomes emphasizes the importance of rodents as hepacivirus hosts and we identify 13 rodent species and 3 rodent genera (in Cricetidae and Muridae families) as novel hosts of hepaciviruses. Through co-phylogenetic analyses, we demonstrate that hepacivirus diversity has been affected by cross-species transmission events against the backdrop of detectable signal of virus-host co-divergence in the deep evolutionary history. Using a Bayesian phylogenetic multidimensional scaling approach, we explore the extent to which host relatedness and geographic distances have structured present-day hepacivirus diversity. Our results provide evidence for a substantial structuring of mammalian hepacivirus diversity by host as well as geography, with a somewhat more irregular diffusion process in geographic space. Finally, using a mechanistic model that accounts for substitution saturation, we provide the first formal estimates of the timescale of hepacivirus evolution and estimate the origin of the genus to be about 22 million years ago. Our results offer a comprehensive overview of the micro- and macroevolutionary processes that have shaped hepacivirus diversity and enhance our understanding of the long-term evolution of the *Hepacivirus* genus.

**Significance:** Since the discovery of Hepatitis C virus, the search for animal virus homologues has gained significant traction, opening up new opportunities to study their origins and long-term evolutionary dynamics. Capitalizing on a large-scale screening of wild mammals, and genomic sequencing, we expand the novel rodent host range of hepaciviruses and document further virus diversity. We infer a significant influence of frequent cross-species transmission as well as some signal for virus-host co-divergence, and find comparative host and geographic structure. We also provide the first formal estimates of the timescale of hepaciviruses indicating an origin of about 22 million years ago. Our study offers new insights in hepacivirus evolutionary dynamics with broadly applicable methods that can support future research in virus evolution.

## Introduction

Thanks to the development of viral detection methods, advances in genome sequencing, and the improvement of computational tools, natural animal reservoirs have been identified for a number of key human viruses. Examples include primates as zoonotic sources of HIV-1 and HIV-2 (Hahn et al. 2000; Sharp and Hahn 2011), multimammate rats as natural hosts of Lassa virus (Olayemi et al. 2016; Bonwitt et al. 2017) and bats harboring a broad diversity of SARS-like coronaviruses (Ge et al. 2013). Although the identification of animal reservoirs of recently emerged pathogens is important, the origins of some viruses with a longstanding history in humans, such as smallpox, hepatitis B virus and hepatitis C virus, remain unresolved.

Hepatitis C virus (HCV) was for a long time the sole representative of the *Hepacivirus* genus within the positive-sense single-stranded RNA virus family *Flaviviridae*. This blood-borne pathogen was discovered in 1989 (Choo et al. 1989) and causes both acute and chronic liver disease, leading to liver cirrhosis and hepatocellular carcinoma in severe cases. According to the World Health Organization, at least 58 million people worldwide have been infected by chronic HCV in 2021 and the number is increasing at a rate of about 1.5 million per year (Spera 2022). While the infection burden of HCV is comparable to that of HIV, evidence of the animal source of this important human pathogen remains lacking.

An effort to shed light on hepaciviruses in animals started in 2011, when HCV homologues were identified for the first time in a non-human host (Kapoor et al. 2011). Since then, a wide variety of HCV-like viruses in diverse animal species have been detected, ranging from mammals to birds, reptiles, arthropods and fish species (Hartlage et al. 2016; Cagliani et al. 2019; Bletsa et al. 2021). So far, the most closely related animal homologue of HCV is the equine hepacivirus. However, it is generally assumed that equids are not the prime candidates for the zoonotic source of HCV because equine hepaciviruses are relatively divergent from HCV, they have a comparably lower genetic diversity, and associated with this, a comparatively more recent time to their most recent common ancestor (tMRCA) (Walter et al. 2017). Rodents and bats harbor the greatest hepacivirus genetic heterogeneity and are considered one of the major transmitters of hepaciviruses to other mammalian species (Quan et al. 2013; Pybus and Thézé 2016; Bletsa et al. 2021). Despite relatively extensive sampling and screening, current efforts have not yet led to a definitive identification of the zoonotic origin of HCV.

In line with the range of hepacivirus host species, these viruses are also geographically broadly distributed. Non-human hepaciviruses have been recorded in countries across six continents: Asia (Quan et al. 2013; Van Nguyen et al. 2018; Wu et al. 2018; Wu et al. 2021), Africa (Corman et al. 2015; Bletsa et al. 2021), North America (Kapoor et al. 2013; Tomlinson et al. 2019), South America (Schmid et al. 2018; de Souza et al. 2019), Europe (Drexler et al. 2013; Kesäniemi et al. 2019) and Australia (Harvey et al. 2019; Porter et al. 2020). Contrary to endemic HCV genotypes that circulate in specific locations, currently identified non-human hepaciviruses do not appear to be restricted to specific areas. Rodents and bats are mainly endemic to most land regions and consequently hepaciviruses from these hosts have been reported in various locations (Quan et al. 2013; Bletsa et al. 2021). Domesticated animals, such as cattle and equids, demonstrate a complex global geographic distribution, mainly resulting from international transport (Walter et al. 2017; Shao et al. 2021; Breitfeld et al. 2022). Apart from these hosts, the majority of hepaciviruses from wild animals tend to exhibit limited spatial ranges, probably due to the restricted habitat range of their hosts. Despite these observations, it is not yet known to what extent diversity within the *Hepacivirus* genus is structured by geography or hosts.

Recent research on the discovery of novel hosts or endogenous viral elements suggests that hepacivirus presence could trace back to million years ago, thus leading to this massive diversity and relatively high prevalence at present (Bamford et al. 2022; Mifsud et al. 2023). However, the potentially ancient origin of hepaciviruses challenges the molecular clock estimation methods that typically rely on contemporary sampling dates of viruses. These methods ignore the time-dependent rate phenomenon (TDRP) that has been commonly observed for the long-term evolution of rapidly-evolving RNA viruses (Duchêne et al. 2014; Aiewsakun and Katzourakis 2016). Among the models available to tackle this problem, Ghafari et al. recently proposed a new mechanistic model that recapitulates the TDRP in power-law rate decay and was initially applied to HCV genotypes (Ghafari et al. 2021). Using this approach, the most recent common ancestor (tMRCA) of HCV genotypes was estimated to date back to about 423 thousand years ago (KYA), so before the modern human movement out-of-Africa. This either supports a single ancient zoonotic origin of HCV and subsequent diversification within modern humans from Africa, or it could be explained by long-term circulation in animal hosts, for which no descendants have currently been sampled, followed by multiple more recent cross species transmissions to humans. A clear timescale for the non-human hepaciviruses and the entire *Hepacivirus* genus is still lacking.

In this study, we screen a comprehensive set of wild animal specimens (*n* = 1,672) originating from more than 75 species mainly from Africa and Asia to characterize the hepacivirus diversity in animal reservoirs. Using available and novel hepacivirus genomes we reconstruct the evolutionary history of hepaciviruses and characterize virus-host co-phylogenetic relationships. Furthermore, we evaluate the influence of host species diversity and geographic distribution on the current phylogenetic structure of hepaciviruses by applying a phylogenetic Bayesian multidimensional scaling (BMDS) approach. Finally, we estimate hepacivirus divergence times and provide a new perspective on the evolutionary timescale of these viruses.

## Material and methods

### Sample collection and hepacivirus detection

To screen for hepacivirus presence in various mammals, a total of 1,672 mammalian samples were collected from several ecological and evolutionary studies (Těšíková et al. 2017). Our sample set contained 1,601 wild small mammals (including shrews, hedgehogs, moles, rodents and bats) from locations in Africa, Asia and Europe. Of these, 214 animals originated from the Democratic Republic of the Congo (DRC), 70 from Guinea, 156 from Senegal, 3 from Tanzania, 713 from China, 64 from Malaysia, 380 from Saudi Arabia and 1 from the Czech Republic (Supplementary Table 1). In addition to our small-mammal collection, a few large-sized mammals were sampled, including 1 civet and 5 galagos from the DRC, and 65 camels from Saudi Arabia.

Whole blood specimens were collected on Serobuvard pre-punched filter papers, which were shipped and stored at room temperature, while tissue specimens, including liver, spleen, kidneys, and muscles, were shipped and stored in RNAlater (Qiagen) at -20°C or in ethanol at room temperature.

Prior to hepacivirus screening, viral RNA was purified from its starting material and reverse transcribed. For 381 samples, RNA extracts were directly provided by our collaborators, while for the remaining 1,291 samples, RNA was extracted in-house. From the latter, RNA was purified for a subset of 220 dried blood spots (DBS) samples (all from DRC) using the QIAamp® Viral RNA Mini kit with a slightly adapted protocol. Briefly, in this modified version, two dried blood spots were used for each sample and incubated for 15 min with 400 μl of AVL-carrier RNA buffer. Upon incubation, we used sterile DNase- and RNase-free micropestles, to extract the blood from the serobuvard filter papers. This process was repeated twice to increase the extraction efficiency. After the second incubation, filter papers were removed and 400 μl of 100% ethanol were added in the tube. For the two washing steps we used a volume of 400 μl of AW1 and AW2, respectively, and final elution was performed with 25 μl of nuclease-free water. For the remaining 1,071 tissue samples, total RNA was purified using the Qiagen RNeasy Mini kit, following the protocol described by Bletsa et al. (2021). This method essentially includes an optimized intermediate on-column DNase treatment to increase RNA yield and purity.

To screen for hepaciviruses, we followed a previously described protocol (Bletsa et al. 2021). In brief, this involved a step of reverse-transcription using Maxima Reverse Transcriptase (ThermoFisher Scientific), random hexamers (ThermoFisher Scientific) and 8 µl of total RNA. Upon generating the complementary DNA (cDNA), a hemi-nested PCR assay was employed using two pairs of degenerate primers that targeted a 300nt fragment of the conserved NS3 protease-helicase genomic region.

PCR products were verified on a 2% agarose gel electrophoresis and the expected products were subsequently purified using ExoSAP-IT PCR Product Cleanup Reagent (Applied Biosystems) or Zymoclean Gel DNA Recovery Kit (Zymo Research). Finally, purified products were sent for Sanger sequencing (Eurogentec, Belgium) and upon delivery of results, a tBLASTx similarity search against a custom hepacivirus-enriched database was used for the detection of hepacivirus positive hits.

### Hepacivirus whole-genome sequencing

Full-length hepacivirus genomes were generated from hepacivirus-positive tissue samples using a meta-transcriptomics approach. Total RNA was first quantified with the Qubit RNA BR assay kit (ThermoFisher Scientific) and qualified using an Agilent RNA 6000 Nano chip (Agilent) in the 2100 Bioanalyzer System (Agilent). Samples with a total RNA Integrity Number (RIN) > 2 were selected for downstream processing.

To increase the proportion of viral RNA reads, ribosomal RNA (rRNA) was depleted prior to library preparation with the NEBNext rRNA Depletion Kit (Human/Mouse/Rat) (Li et al. 2022, DOI: dx.doi.org/10.17504/protocols.io.q26g74843gwz/v1). Before subjecting all samples to this rRNA depletion step, we assessed hepaciviral RNA yield using a custom real-time quantitative PCR (qPCR) assay. This qPCR assay was performed with the Universal KAPA SYBR FAST qPCR kit (Sigma Aldrick) on the ABI 7500 Fast Real-Time PCR desktop system (Applied Biosystems). In brief, cDNA was generated using Maxima Reverse Transcriptase (ThermoFisher Scientific) with random hexamers (ThermoFisher Scientific) following the manufacturer’s protocol. The internal primer pair AK4340F2 and AK4630R2 from our hemi-nested screening PCR was used for quantification of hepacivirus RNA. Standard curves were calculated based on serial dilutions of a 393bp fragment of the NS3 genomic region from one of our hepaci-positive samples (sample voucher: S2645, NCBI accession no. OM162002). To assess mitochondrial rRNA depletion, we designed two pan-rodent primers 12S-512F and 12S-694R (Supplementary Table 2) targeting a ∼183bp fragment of the 12S rRNA rodent mitochondrial genomic region. Standard curves were calculated using serial dilutions of the aforementioned gene fragment from a *Lophyromys* mice (NCBI accession no. AJ250349). The qPCR assay was performed with the Universal KAPA SYBR FAST qPCR kit (Sigma Aldrich) on an Applied Biosystems 7500 Fast Real-Time PCR desktop system (Applied Biosystems) using the following cycling conditions: enzyme activation at 95 °C for 3 min; 40 cycles of denaturation at 95 °C for 3s, annealing/extension at 57 °C for 30 s (for the NS3 hepacivirus), while 58 °C in 30 s (for the 12S rRNA); (3) dissociation using the default settings of our qPCR platform. All qPCR data were analyzed with ABI PRISM 7000 software (v2.0.5; Applied Biosystems).

Upon rRNA depletion, the quantity and quality of total RNA were assessed using the Qubit RNA HS assay kit (ThermoFisher Scientific) and the Agilent RNA 6000 Pico kit (Agilent), respectively. Sequencing libraries were prepared using the NextFlex Rapid directional RNAseq kit (PerkinElmer). RNA input volume was determined by the total amount of ∼100 ng and adjusted to 14 µl with nuclease-free water. For the library preparation assay we generally followed the manufacturer’s guidelines with some optimizations in two steps (Li et al. 2022, DOI: dx.doi.org/10.17504/protocols.io.kqdg3p6pel25/v1). Specifically, we adjusted the incubation time for scalable RNA fragmentation and the number of PCR amplification cycles were determined in a sample-specific fashion depending on the quality and amount of starting material.

NGS libraries were subsequently checked for quality and quantified using the Agilent High Sensitivity DNA kit and the KAPA Library Quantification Kit Complete Kit (ROX Low) for Illumina Platforms, following the manufacturer’s protocol. Libraries were normalized using Tris-HCl (10 mM) with 0.1% Tween 20, followed by pooling and paired-end high-throughput sequencing on an Illumina NextSeq 500 platform at the VIB Nucleomics Core (Flanders, Belgium).

### Hepacivirus genome assembly and host species identification

We built on a previously described high-throughput sequencing data analysis pipeline to assemble complete hepacivirus genomes (Bletsa et al. 2021) and further optimized it to increase its efficiency. Demultiplexing was performed using Bcl2fastq2 v2.20 (Illumina). Quality of the raw read data was checked in FastQC v0.11.9 (Andrews 2010), followed by quality filtering and Illumina adaptor trimming using Trimmomatic v0.39 (Bolger et al. 2014). To subtract host background, a total number of 11 genomes from various rodent species and a human genome (NCBI accession no. in supplementary table 3) were used as mapping references in SNAP aligner v1.0.3 (Zaharia et al. 2011). PRINSEQ-lite v0.20.4 (Schmieder and Edwards 2011) was used to filter out duplicates and low complexity reads. A *de novo* assembly approach was followed to generate contigs using SPAdes genome assembler v3.15.2 (Bankevich et al. 2012). All generated contigs were screened using the BLASTx algorithm against a custom hepacivirus-enriched viral protein database in DIAMOND v2.0.9 (Buchfink et al. 2021). To increase the overlap between all identified hepacivirus contigs or create scaffolds from discontinuous contigs, re-assembly was performed in CAP3 version date 02/10/15 (Huang and Madan 1999). Coverage and sequencing depth statistics were calculated by remapping all pre-processed reads to each newly assembled hepacivirus genome with Bowtie2 v2.4.2 (Langmead and Salzberg 2012), coupled with SAMtools v1.12 (Danecek et al. 2021) for format conversion and visualized using the weeSAM script (https://github.com/centre-for-virus-research/weeSAM).

For several samples resulting in partial hepacivirus genomes, strain-specific PCR assays were designed to fill genomic gaps (Supplementary Table 2). Overlapping amplicons were generated using the OneStep RT-PCR kit (Qiagen) with 5 μl of cDNA as a template. PCR products were purified and Sanger sequenced in both directions. Sequences were mapped and concatenated to their original contigs in Geneious Prime v2020.2.4 (Biotmatters, Auckland, New Zealand, https://www.geneious.com) to obtain complete viral genomes.

For the purpose of host species identification, the mitochondrial cytochrome b (cytb) gene was reconstructed by *de novo* assembling the trimmed read data. Upon contig generation, a total number of 52,405 rodent cytb sequences were downloaded from the NCBI database (search of 2021 September 27th) for building a local BLAST reference database in BLASTn v2.10.1+ (Camacho et al. 2009). The tBLASTx algorithm along with a custom perl script were used to extract cytb contigs, followed by similarity search against the NCBI nt database. To delineate any unclassified rodent species we relied on our collaborators’ expert opinion.

### Phylogenetic analysis

All hepacivirus nucleotide sequences generated were first translated to amino acid using Aliview v1.27 (Larsson 2014). The polyprotein coding regions were predicted based on all available rodent hepacivirus complete genomes from NCBI.

To reconstruct the phylogeny of the whole *Hepacivirus* genus, we analyzed our novel hepacivirus genomes (*n* = 34) along with all available complete hepacivirus polyprotein sequences (*n* = 259, search on August 2022) with annotated host species information. The latter were downloaded from NCBI and included 21 hepaciviruses from cattle, 1 from dog, 48 from equids, 7 of human origin (one representative reference genome per genotype), 10 of non-human primate origin, 2 hepaciviruses from sloths, 2 from ringtails, 2 from marsupials, 125 from rodents, 11 from bats, 3 from shrews, and 27 from non-mammalian hosts (Supplementary table 4). For the human HCV subset, in addition to including one representative reference genome per genotype, we also collected specific subsets for genotypes 1a (*n* = 35), 1b (*n* = 34) and 3a (*n* = 35) for further downstream evolutionary analyses.

The complete amino acid dataset was aligned using MAFFT v7.453 (Katoh 2002) in a stepwise approach. In brief, sequences from the same hosts were initially aligned in batches and we progressively incorporated the various host-specific alignments into a single Multiple Sequence Alignment (MSA). All alignments were visually assessed and manually edited using Aliview v1.27 (Larsson 2014). The phylogenetic informative blocks were selected using a BLOSUM30 matrix and trimmed with the gap frequency criteria of 0.7 using BMGE v1.12 (Criscuolo and Gribaldo 2010). Upon obtaining our complete MSA, IQ-TREE v1.6.12 (Nguyen et al. 2015) was used to reconstruct the hepacivirus phylogeny with 1000 bootstrap replicates. The best-fitting amino acid substitution model according to BIC was LG+F+I+G4. Trees were visualized and annotated in Figtree v1.4.4 (http://tree.bio.ed.ac.uk/software/figtree/).

To further explore the evolutionary relationships among different hepaciviruses, the conserved regions in NS3 (position 1123 – 1566 in amino acid, with AAA45676 as reference) and NS5B (position 2536 – 2959 in amino acid), which have been previously used for species classification of members of the *Hepacivirus* genus (Smith et al. 2016), were extracted from the complete genome. The amino acid alignments were used to compute pairwise p-distance in MEGA11 (Tamura et al. 2021). The corresponding genetic distances heatmaps were generated using ComplexHeatmap R package (Gu et al. 2016).

### Virus-host co-divergence analysis

To assess co-phylogenetic relationships between hepaciviruses and their hosts, we compared the topological structure of virus and host phylogenies. For these analyses, we downsampled our genome-wide hepacivirus dataset by retaining only one representative hepacivirus genome per host species from the same lineage. In addition, the basal hepacivirus sequences originating from non-mammalian hosts were removed due to the high degree of genetic divergence and the uncertainty of host species in the case of blood-feeding arthropods. This resulted in a final hepacivirus dataset of 85 sequences belonging to 58 mammalian taxa. To reconstruct the mammalian host species phylogeny, we extracted a 31-gene supermatrix alignment for a subset of 51 host taxa based on the most comprehensive mammalian species classification to date (Upham et al. 2019). For the remaining 7 host species, which were not included in the 31-gene collection, we manually added mitochondrial cytb sequences to the supermatrix alignment (Supplementary Table 5) The final MSA was generated using MAFFT v7.453 (Katoh 2002), manually edited in Aliview (Larsson 2014) and a phylogenetic tree was constructed with IQ-TREE v1.6.12 (Nguyen et al. 2015) using the best BIC fitting model GTR+F+I+G4.

We used the event-based eMPRess software (Santichaivekin et al. 2021) and the Procrustes global-fit test PACo (Balbuena et al. 2013) to assess phylogenetic congruence between the host and hepacivirus trees. The significance of the single viral-host link was evaluated using AxParafit (Stamatakis et al. 2007) through Copycat (Meier-Kolthoff et al. 2007). In the eMPRess analysis, upon determining the most parsimonious costs for duplications, transfers and losses, we performed a permutation test with 100 randomizations to calculate the support value. For PACo and AxParafit settings, the co-phylogenetic signals were evaluated based on 100,000 permutations and considered to be significant if the observed squared residual distance was smaller than 0.05. Visual correspondence (tanglegrams) between the two phylogenies was created using the phytools v1.2-0 (Revell 2012) R package.

Due to the high proportion of hepacivirus co-infections in *Lophuromys* mice, we performed a second virus-host co-divergence analysis on a reduced dataset. In this latter dataset, we removed all hepacivirus sequences that were retrieved from co-infected mice and only retained genomic information from single hepacivirus infections (*n* = 63) and their corresponding host species (*n* = 57). For the virus-host co-phylo plot and the phylogenetic congruence analysis we followed the approach described above.

### Evaluating hepacivirus geographic versus host structure

As a first step towards exploring the phylogeographic clustering of hepaciviruses, we compiled sampling location information for all the sequences in our genome-wide dataset and collected the corresponding geographic coordinates. When only country information was available, we used the geographic coordinates from the capital city of the sampled country. To visualize the spatial distribution of hepacivirus, a country-level allocation plot was generated with ggtree v3.2.1 (Yu et al. 2017) and ggplot2 v3.3.5 (Wickham 2009) in R.

In order to formally compare to what extent hepaciviruses are structured by geography or host we explored a new approach based on phylogenetic Bayesian multidimensional scaling (BMDS) (Bedford et al. 2014). Our approach sidesteps the problem of discrete approaches that require arbitrary discretizations according to geography and hosts, which are highly likely to differ in their dimensionality and hence difficult to compare in terms of their phylogenetic association. Our BMDS approach conditions on geographic distances and host distances between pairs of hepaciviruses and estimated locations in a lower dimensional geographic and host space for the sampled viruses (phylogenetic tips) and their hypothetical common ancestors (internal nodes). In our probabilistic BMDS formulation, observed distances are assumed to be centered on their mapped distances with a Gaussian error. As a prior on the unobserved locations in lower-dimensional space, we specify a phylogenetic Brownian diffusion process (Lemey et al. 2010). To determine the appropriate dimensionality for our BMDS procedure, we adopted a cross-validation approach (Holbrook et al. 2021).

We proposed two metrics to compare the diffusion process in geographic and host space. One of these metrics is Pagel’s lambda parameter (Pagel 1999), which measures the degree of phylogenetic association of a continuously-valued trait. A Pagel’s lambda estimate close to 0 reflects the absence of phylogenetic association whereas an estimate close to 1 reflects a phylogenetic signal that is expected under a Brownian diffusion process. The second metric is the standard deviation of a relaxed random walk (RRW) process, which relaxes the constant-variance assumption of Brownian diffusion. This metric reflects the regularity of diffusion in geographic or host space, with a higher standard deviation representing a more heterogeneous diffusion process. We used a Bayesian implementation to co-estimate Pagel’s lambda in our BMDS approach (Vrancken et al. 2015) and performed two analyses. The first analysis models standard independent Brownian diffusion processes in lower dimensional geographic and host space on a random phylogeny, incorporating a Pagel’s lambda parameter for both diffusion processes. In this analysis we estimated the phylogeny using a LG amino acid substitution model with a discrete gamma distribution to model among-site rate variation and a Yule speciation prior. We specified an uncorrelated relaxed molecular clock model with mean fixed to 1 (so, estimating branch lengths in substitution units). The second analysis models independent RRW diffusion processes in both geographic and host space using a lognormal distribution with an estimable standard deviation on a fixed tree topology while also incorporating the Pagel’s lambda estimator. As a fixed tree, we used the MCC tree (or relevant subtree) rescaled by the time-dependent rate modelling in our dating approach (cfr. Methods, section 2.8). We applied both analyses to all non-mammalian hepaciviruses (*n* = 13), all mammalian hepaciviruses (*n* = 257), a mammalian subset with only a single representative for clusters of viruses that were sampled from the same host and country (*n* = 160), a subset of the latter with only a single representative for bovine and equine viruses (*n* = 123), only rodent viruses (*n* = 95) and only rodent viruses excluding those sampled in co-infections (*n* = 68).

The mammalian host phylogeny that served as the basis for the host distances (*n* = 58 host taxa) was generated according to the description in Methods (section 2.5). To compile the non-mammalian dataset (*n* = 13 host taxa), we selected from the genome-wide hepacivirus dataset the non-mammalian host species for which we had geographic information. For those host species, we visited the VertLife.org (http://vertlife.org) database and inferred separate phylogenies for the fish (*n* = 4 species), the squamates (*n* = 3 species) and the birds (*n* = 4 species) in our dataset. Based on the alignments that gave rise to a relatively recent and robust vertebrate phylogeny (Irisarri et al. 2017), we created a backbone tree with 5 taxa (in which 3 taxa originated from host species in our vertebrate hepacivirus dataset and 2 species were used as calibrations for the merging of the subtrees) (Supplementary Table 6). Upon generating all subtrees, we used the tree.merger R package (Castiglione et al. 2022) to incorporate the subtrees into the backbone vertebrate phylogeny in a stepwise fashion. Host distances were extracted as patristic distances from both the mammalian and the non-mammalian phylogeny using R and geographic distance matrices were constructed using great-circle distances based on the coordinates (cfr. previous section).

### Recombination and temporal signal analysis

To avoid the impact of recombination prior to assessing temporal signal and inferring time-scaled phylogenies (Schierup and Hein 2000; Arenas and Posada 2010; Martin et al. 2011), we performed recombination analyses on a restricted number of host-specific hepacivirus lineages with comparatively low phylogenetic diversity. For these analyses, we selected the entire collection of cattle (*n* = 21), equids (*n* = 48), HCV genotype 1a (*n* = 35), HCV genotype 1b (*n* = 34), HCV genotype 3a (*n* = 35) and three rodent hepacivirus lineages (*n* = 13, *n* = 25, and *n* = 50) (Supplementary Fig. 1).

For those different subsets, nucleotide sequences of the complete polyprotein were aligned as codons using MUSCLE in MEGA11 (Tamura et al. 2021). A recombination evaluation was conducted using the Phi test (Bruen et al. 2006) (window size 100, significance threshold = 0.05) in SplitsTree4 v4.18.2 (Huson and Bryant 2006). To further test for evidence of recombination, we employed 7 different detection algorithms in the RDP4 program (Martin et al. 2015) under the following conditions: RDP (window size 30), BootScan (window size 200, step size 20), SiScan (window size 200, step size 20), GENECONV, Chimaera, MaxChi, and 3Seq (default settings). The highest acceptable p-value was set at 0.05 and recombination events were reported only upon detection by more than 3 methods. Recombinant-free alignments were generated by masking the minor recombinant regions.

To examine the relationship between genetic diversity and sampling date, we estimated temporal signal using the non-recombinant regions of the lineage-specific alignments. Sequences without known sampling date were removed, which resulted in the final subsets of cattle (*n* = 19), equids (*n* = 35), HCV genotype 1a (*n* = 35), HCV genotype 1b (*n* = 34), HCV genotype 3a (*n* = 35), rodent lineage 1 (*n* = 13), lineage 2 (*n* = 25) and lineage 3 (*n* = 50). We used TempEst v1.5.3 (Rambaut et al. 2016) for a visual inspection of the dataset by plotting the root-to-tip divergences against sampling time based on the non-molecular clock tree built using IQ-TREE v1.6.12 (Nguyen et al. 2015). A more formal analysis was performed in BEAST v1.10.4 (Suchard et al. 2018) with the high-performance BEAGLE phylogenetic compute library (Ayres et al. 2012) using generalized stepping-stone sampling (GSS) (Baele et al. 2016) marginal likelihood estimation with an initial Markov chain length of 200 million and 50 stepping stones each with a chain length of 1 million.

### Divergence time estimation using the PoW model

To estimate a timescale for hepacivirus evolutionary history, we applied the ‘Prisoner of War’ (PoW) evolutionary rate decay model developed by Ghafari et al. (2021). This model takes into account that sequence divergence is impacted by substitution saturation and following a recent period of evolution at the short-term evolutionary rate, lower evolutionary rates apply to deeper parts of the evolutionary history following a universal rate decay dynamic.

The PoW model requires an estimate of the short-term hepacivirus evolutionary rate. In the absence of temporal signal in most hepacivirus lineages, we estimated this rate using a dated tip model applied to HCV genomes with sampling years ranging between 1990 and 2015. In order to obtain a rate estimate for the exact same sites as in the hepacivirus data set, we first combined HCV genotype 1a, 1b and 3a sequences (*n* = 104) with our previously described hepacivirus dataset (*n* = 293). Multiple sequence alignment was performed at the amino acid level using MAFFT v7.453 (Katoh 2002) and the conserved regions were selected using BMGE v1.12 (Criscuolo and Gribaldo 2010) with 20 block size and BLOSUM75 matrix. In particular, from an initial alignment that spanned 6,027 amino acid residues we extracted an alignment of 669 relatively conserved amino acid sites. The conserved alignment was back translated to nucleotides in TranslatorX (Abascal et al. 2010), and separated into the HCV genotype 1a, 1b and 3a dataset on the one hand and the hepacivirus data set on the other hand. Using the Phi test, we did not find any detectable recombination in this hepacivirus data set (p = 0.98). We used a strict clock model with dated tips applied to the three independent phylogenies of HCV1a, 1b and 3a and HKY substitution model to estimate the short-term rate in BEAST v1.10.4 (Suchard et al. 2018) with BEAGLE (Ayres et al. 2012) using an MCMC of 200 million generations and a 10% burn-in. We also used BEAST to estimate a posterior distribution of trees in units of genetic distance for the hepacivirus data set specifying again an HKY substitution model. Using the short-term evolutionary estimate and adopting the previously predicted maximum substitution rate for single-stranded RNA (ssRNA) viruses of 3.65×10^-2^ substitutions per site per year (Ghafari et al. 2021), the hepacivirus tree distribution was transformed to units of time under the PoW model.

## Results

### Novel hepaciviruses present in a wide range of rodent species

We screened for hepaciviruses a comprehensive set of wild mammal samples (*n* = 1,672) that were obtained by our collaborators in Africa, Asia and Europe. The majority of our samples were small mammals (*n* = 1,601) that belong to 75 potential species across 47 genera. Specifically, 169 specimens belong to 14 Insectivora species (Order: Eulipotyphla), 1,396 specimens originated from at least 50 rodent species (Order: Rodentia) and 36 specimens were collected from 8 bat species (Order: Chiroptera). In addition to the small-mammal samples, we also included specimens from 1 civet, 5 galagoes, and 65 camels in our screening efforts (Supplementary Table 1).

After molecular detection, a total of 53 hepacivirus positive specimens were identified across 18 host species within the Rodentia order. Out of those 18 host species, 14 species belong to the Muridae family and 4 species to the Cricetidae family (Supplementary Fig. 2 A). In the Muridae family, we extend the hepacivirus host repertoire by identifying 10 new hepacivirus-positive rodent species that belong to 7 genera: *Acomys*, *Dipodillus*, *Lophuromys*, *Meriones*, *Micromys*, *Niviventer*, and *Rattus*. Among them, the *Dipodillus* and *Micromys* genera were for the first time identified as hepacivirus hosts in this study (Supplementary Table 7). In the Cricetidae family, we identified 3 novel hepacivirus-positive host species from the *Eothenomys* genus, and provided additional proof that rodents from the *Clethrionomys* genus harbor hepaciviruses. Despite previous evidence that shrews and bats can carry hepaciviruses, we were not able to detect any hepaciviruses from those animals, but we only had limited samples available for these hosts. In addition, none of the very few surveyed galagoes, and camels tested positive for hepaciviruses, nor did the civet.

With respect to the percentage of positive rodent specimens in this study, we detected the lowest hepacivirus prevalence (0.88%, 1 out of 114) in *Niviventer confucianus*, followed by *Rattus rattus* with 1.49% (1 out of 67) and *Praomys* mice with 2.94% (1 out of 34). While sampling sizes were considerably smaller for species of the *Eothenomys* genus compared to the previously mentioned species, they exhibited much higher prevalence. In particular, the positivity rate for *Eothenomys eleusis* was 55.56% (5 out of 9), for *Eothenomys miletus* was 40% (2 out of 5) and for *Eothenomys melanongaster* was 22.22% (4 out of 18). In line with the high prevalence in *Eothenomys* species, the positivity rate of *Rattus andamanensis* was 50% (1 out of 2), while the hepacivirus-positive percentages for the remaining species ranged from 2.94% to 26.32% (Supplementary Table 1).

As for the spatial distribution of our novel hepaciviruses, the majority were detected in Saudi Arabia with a proportion of 7.37%, followed by Guinea with 4.29%. Rodents from China and the DRC presented a similar hepacivirus positive rate, with 1.96% and 1.87% respectively. Hepaciviruses in a bank vole from the Czech Republic and in three Tanzanian samples were detected during a metagenomic sequencing effort and were not considered in the prevalence calculations, since no hepacivirus-targeted molecular screening was performed. For a more detailed summary of our screening results by country and mammalian species, see Supplementary Tables 1 and 7, respectively.

### Phylogenetic reconstruction demonstrates high hepacivirus diversity

We generated 34 complete hepacivirus genomes from 25 samples belonging to 13 rodent host species. Our results contribute significant new genomic data to the currently available rodent hepacivirus data, mainly by adding to the genomes available from China (14 new) and Tanzania (8 new). In addition, we provide novel genomic records from three previously unrepresented locations: Saudi Arabia (4 new), Czech Republic (1 new) and Guinea (7 new) (Supplementary Fig. 3).

Through genome-wide phylogenetic reconstruction, we confirm the high genetic heterogeneity of the *Hepacivirus* genus (Fig. 1). In agreement with Bletsa et al. (2021), mammalian hosts harbor highly divergent viruses, with only the bovine, equine-canine, marsupial, sloth, ringtail and human hepaciviruses clustering into monophyletic clades. Viruses from rodent, bat and non-human primate species on the other hand are interspersed throughout the phylogenetic tree, forming multiple divergent lineages and exhibiting the largest mammalian hepacivirus diversity. The rodent and bat groups host the majority of assigned species within the *Hepacivirus* genus at present.

**Fig. 1.**
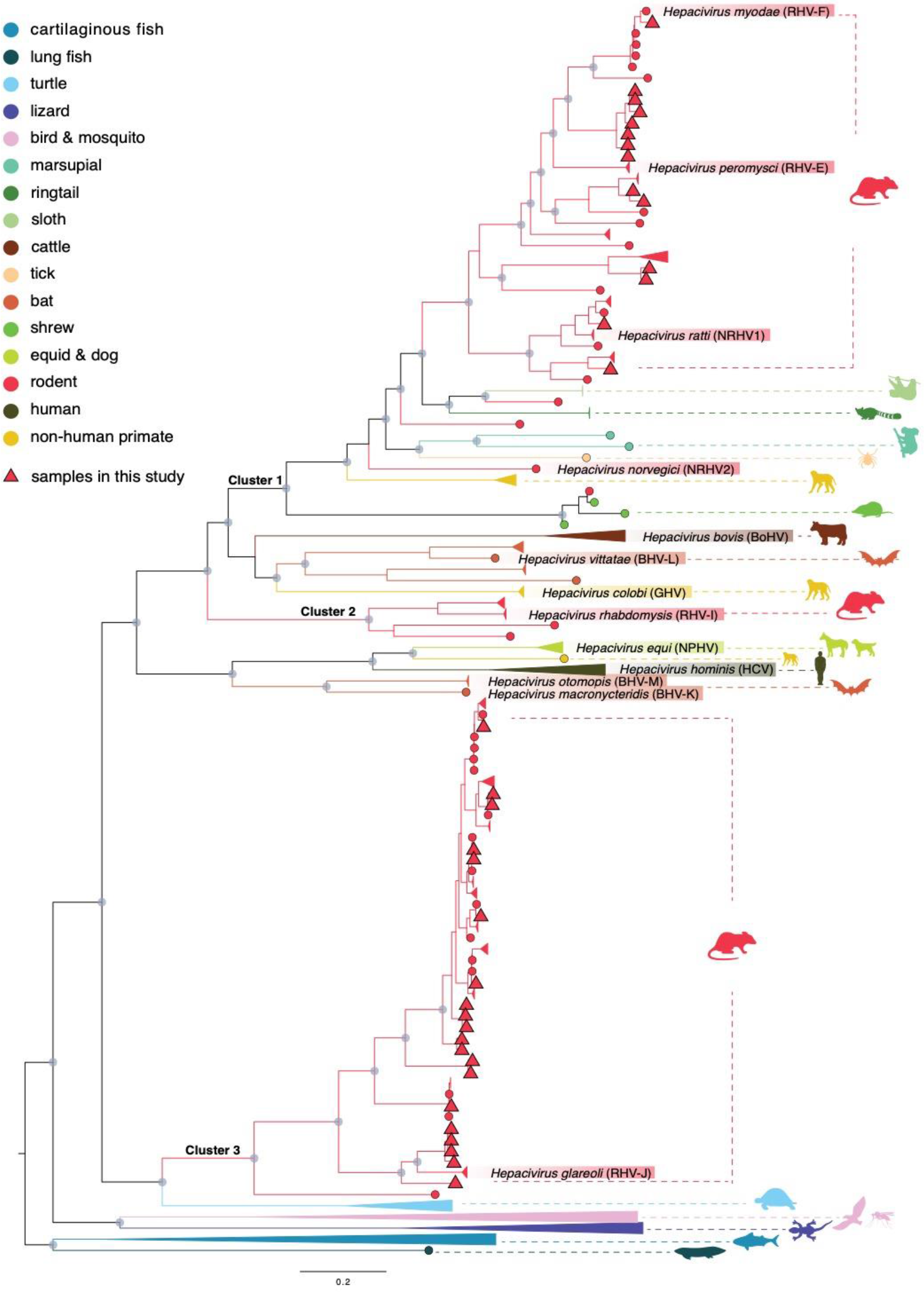
Phylogenetic reconstruction of hepaciviruses based on complete genomes. Selected clades have been collapsed to emphasize the rodent hepacivirus relationships. Tips in circles indicate sequences generated in previous studies (*n* = 259). Tips in triangles indicate novel sequences generated in this study (n = 34). Clades are colored based on the host type as represented in legend. Internal nodes with Shimodaira-Hasegawa (SH)-like support values ≥ 80 are labeled with gray circles. The scale bar indicates the number of amino acid substitutions per site. To better frame phylogenetic relationships, the current demarcations of hepacivirus species and their abbreviations by ICTV are highlighted in coloured boxes at the tips of the trees. The 3 major clusters containing rodent hepaciviruses are labeled for future reference (clusters 1-3).

All our novel rodent hepaciviruses (RHVs) fall into 2 major rodent clusters, denoted as cluster 1 and cluster 3 in Fig. 1. The former is positioned within the heterogeneous mammalian hepacivirus group, while the latter is composed solely of rodent hepaciviruses (Fig. 1 & 2). To illustrate better the phylogenetic relationships of our novel hepaciviruses within the overall rodent hepaciviruses diversity, we computed amino acid genetic distances for all rodent hepaciviruses of cluster 1 and cluster 3 based on the conserved NS3 and NS5B regions according to Smith et al. (2016).

**Fig. 2.**
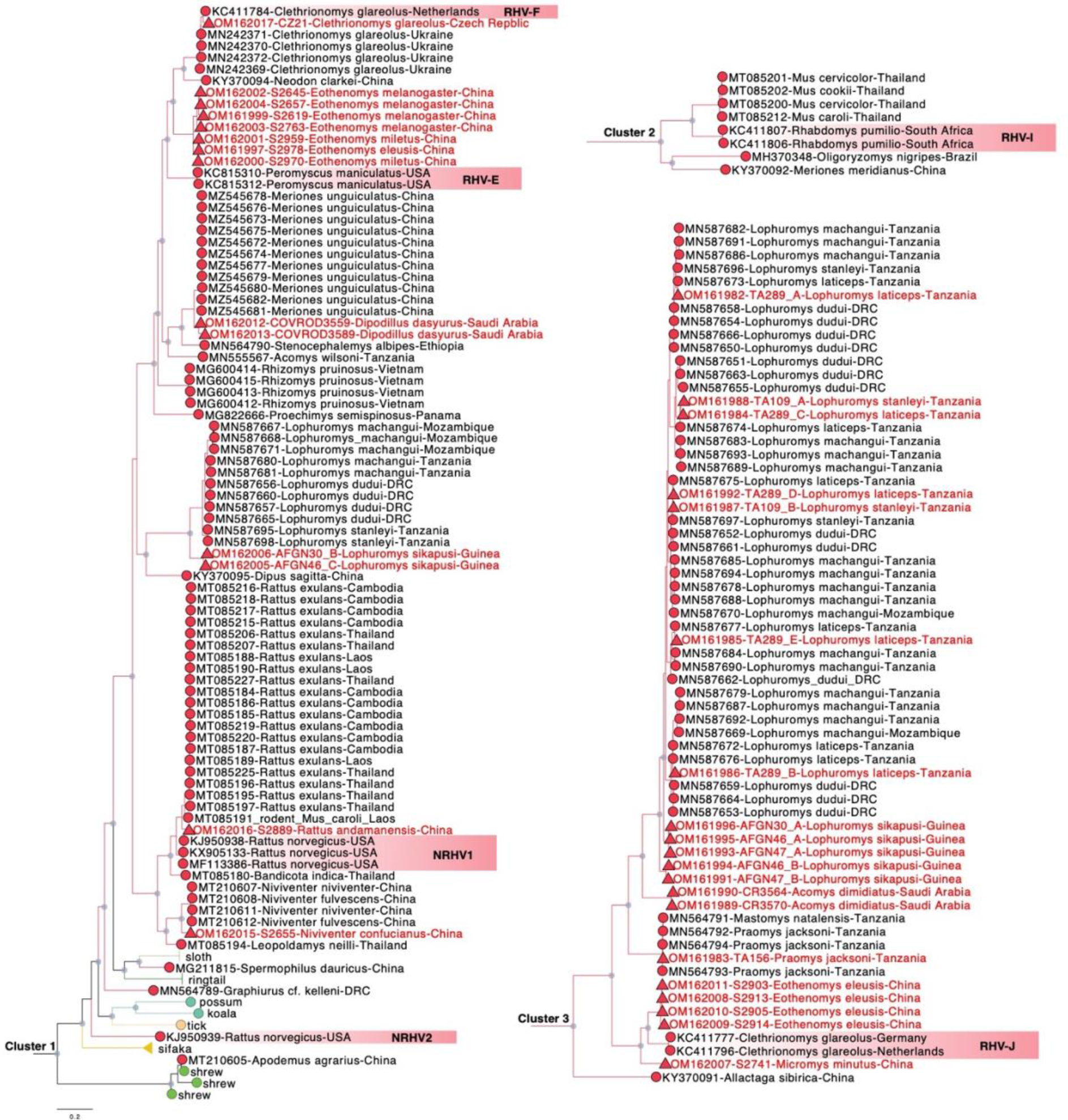
Phylogenetic grouping of rodent hepaciviruses within their three major clusters. Tips with red circles indicate the rodent sequences (*n=* 125) generated in previous studies. Tips with triangles indicate the novel rodent hepacivirus sequences (*n* = 34) generated in this study. Internal nodes with bootstrap values ≥ 80 are labeled with gray circles. The scale bar indicates the number of amino acid substitutions per site. The ICTV classified hepacivirus species are highlighted in coloured boxes at the tips of the tree as shown in Fig. 1.

As shown in Fig. 2, the new RHV genomes from *Clethrionomys glareolus* and *Praomys jacksoni* cluster within the clades of previously identified RHVs from those same rodent species. The estimated amino acid genetic distances for those viruses are < 0.07 in both the NS3 and NS5B regions (Suppl. Figure 4). Hepaciviruses from *Rattus andamanensis* and *Niviventer confucianus* individuals are very closely related to hepaciviruses from other rodent species of the *Rattus* and *Niviventer* genera, respectively (Fig. 2). Amino acid genetic distances in these cases were computed to be < 0.15 in both the NS3 and NS5B regions. *Dipodillus* rodents are newly identified hepacivirus hosts and they harbor RHVs related to those from the *Meriones* genus. Both the *Dipodillus* and the *Meriones* genera belong to the Gerbillinae subfamily; in these rodents we observe a clustering of hepaciviruses at the rodent subfamily level.

Despite the non-random clustering of hepaciviruses from the aforementioned rodent taxa, there are RHVs that do not closely follow host relatedness. Specifically, the Chinese *Micromys minutus* hepacivirus groups with viruses sampled from European *Clethrionomys* and Chinese *Eothenomys* genera. Interestingly, the latter genus even belongs to a different rodent family (Cricetidae) than the *Micromys* rodents (Muridae). Another striking example is the clustering of our newly discovered *Acomys dimidiatus* hepaciviruses from Saudi Arabia. These strains group with viruses obtained from Guinean *Lophuromys* mice and are very divergent from the hepacivirus found in a Tanzanian *Acomys wilsoni* individual (Fig. 2).

Consistent with the results of Bletsa et al. (2021), we find that *Lophuromys* mice commonly exhibit hepacivirus co-infections. Specifically, in *Lophuromys* rodents from Tanzania we identified between two (specimen TA109) up to five (specimen TA289) hepacivirus strains in the same individual. These sequences form divergent lineages in the monophyletic cluster 3 and group with RHVs from the same host species. In addition to the multiple co-circulating hepaciviruses found in Tanzanian *Lophuromys* mice, we also identified RHV co-infections in two Guinean *Lophuromys sikapusi* individuals. These hepaciviruses have a close phylogenetic relationship with those circulating in central/eastern Africa and demonstrate an amino acid genetic distance < 0.13 in NS3 and < 0.08 in NS5B.

Finally, all hepaciviruses detected in *Eothenomys* species form distinct lineages within cluster 1 and cluster 3. Viruses in cluster 1 exhibit high similarity to the *Hepacivirus peromysci* (RHV-E) species and *Hepacivirus myodae* (RHV-F) species. The amino acid genetic distances between *Eothenomys* hepaciviruses and RHV-E were estimated to range between 0.24 - 0.27 in NS3 and 0.23 - 0.27 in NS5B, while the respective genetic distances between *Eothenomys* hepaciviruses and RHV-F vary between 0.26 - 0.28 in NS3 and 0.22 in NS5B. As for the *Eothenomys* viruses in cluster 3, they form sister lineages with the *Hepacivirus glareoli* (RHV-J) species. For these strains, the amino acid genetic distances between *Eothenomys* hepaciviruses and RHV-J varied between 0.13 – 0.15 in NS3 and was estimated to be 0.13 in the NS5B region.

### Impact of host and geography on hepacivirus diversification

To evaluate the extent to which host relatedness impacts hepacivirus diversification, we measured the cophylogenetic signal between hepaciviruses and their hosts by conducting a reconciliation analysis. Although our results provide evidence for significant cophylogenetic patterns using both the eMPRess event-based test (p-value < 0.01) and the Procrustes global-fit test (PACo p-value < 0.001, AxParafit p < 0.001, using 100000 permutations), the tanglegram visualization also demonstrates numerous cross-species transmission events. Noticeably, the *Lophuromys* mice exhibit several interlaced connections and have a significant contribution to co-speciation events, which may be the result of frequent co-infections found in those species (Fig. 3 & 4). Considering this, we re-ran the reconciliation analysis after removing all the co-infections from our dataset. In this latter analysis, the overall global test results support significant phylogenetic congruence between host and mammalian hepaciviruses in all methods tested (eMPRess p-value < 0.01, PACo p-value < 0.001 and AxParafit p < 0.01) and display the topological structure of the virus and the host phylogenies with low similarity (Supplementary Fig. 5 & 6).

**Fig. 3.**
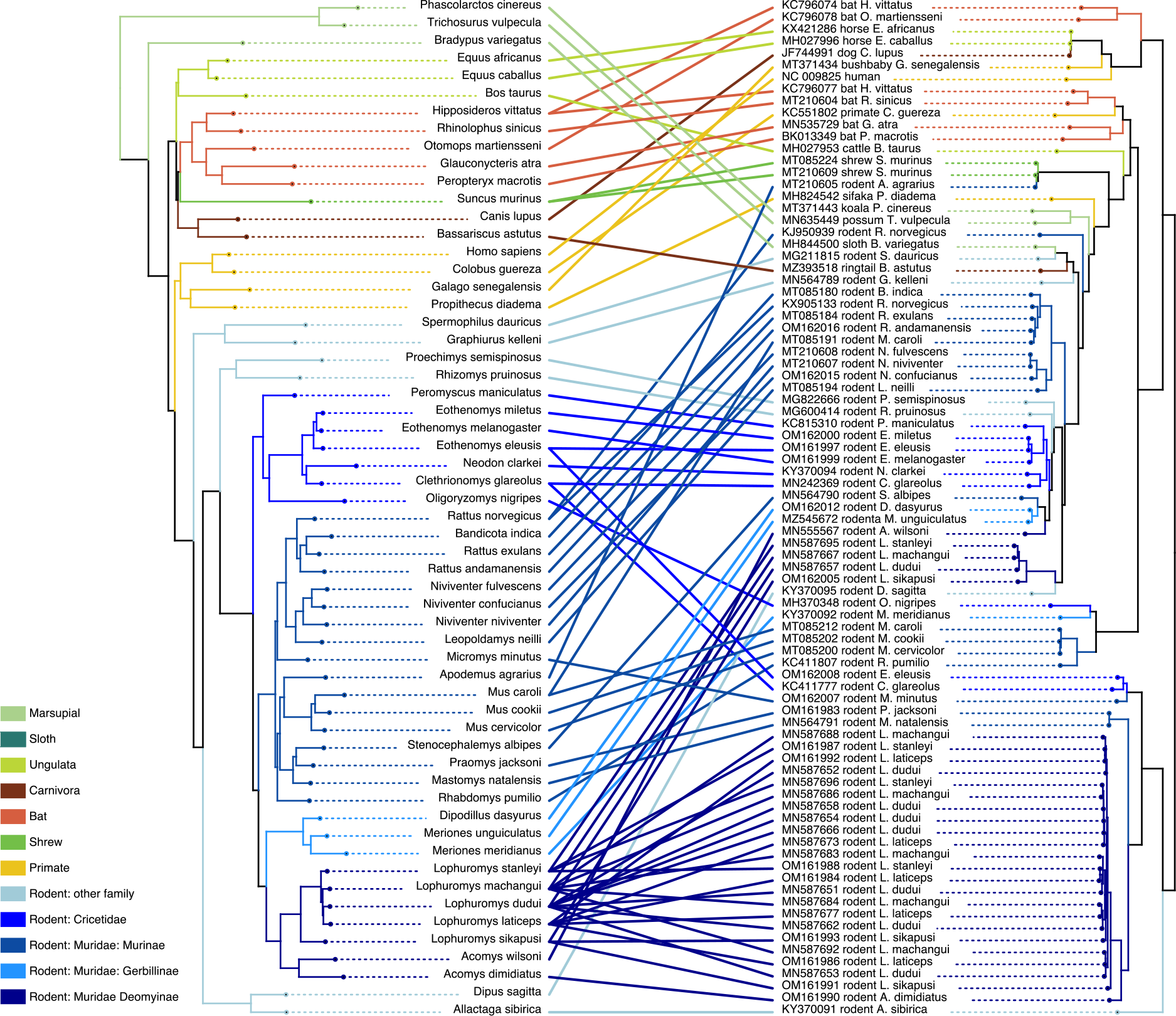
Tanglegram of host (left) and hepaciviruses (right). The host phylogeny was inferred for 31 genes from 58 mammalian species. For *Myodes glareolus*, *Gerbillus dasyurus* and *Microtus clarkei* their updated species names were used: *Clethrionomys glareolus*, *Dipodillus dasyurus* and *Neodon clarkei*, respectively. The hepacivirus phylogeny was inferred for 85 representative genomes. Clades and associations were colored based on host type.

**Fig. 4.**
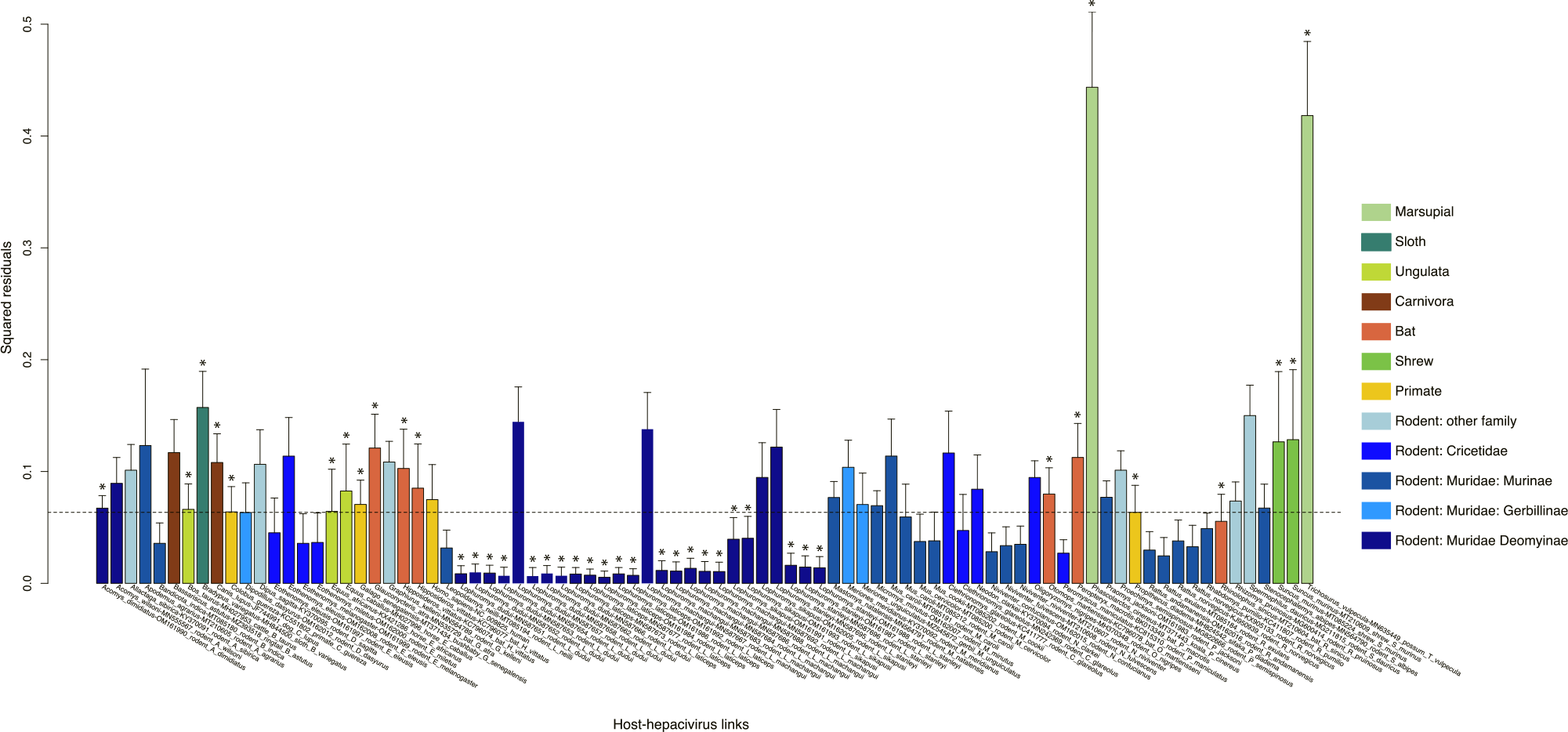
Contribution of each host-hepacivirus association to the general co-evolution pattern. Bars represent Jack-knifed squared residuals with the upper 95% confidence intervals from PACo test. The median squared residual value is shown with a dashed line. Stars represent the host-hepacivirus links that were also tested to be significant in AxParafit (p < 0.05). Colors represent different host types as shown in Fig. 3.

To explore visually the geographic structure of hepacivirus diversity, we colored the viral clades according to their country of sampling and examined their distribution (Fig. 5). Available genomes complemented with our novel data predominantly represent hepaciviruses from Africa and Asia. Due to limited sampling, hepaciviruses from non-mammalian hosts, marsupial, carnivora and shrews are represented by only a restricted number of locations. Equine and bovine hepacivirus clades both contain sequences sampled worldwide, which are relatively intermixed, thus suggesting a dynamic dispersal history likely impacted by anthropogenic factors. Thanks to a relatively extensive sampling, rodent hepaciviruses cover a much broader spatial distribution. In the coinfection clade of *Lophuromys* hepaciviruses we exclusively observe African-specific viruses, while all the other RHVs tend to be structured by location. One such example are RHVs originating from Asia, which form several small clusters and are distinct from those originating from Europe or Africa. In addition, evidence of potential geographic isolation is also observed in bat hepaciviruses, although on a much more limited sampling. In conclusion, we observe that both host species and geography have shaped hepacivirus diversification, however, it remains challenging to determine their relative impact without a formal comparison.

**Fig. 5.**
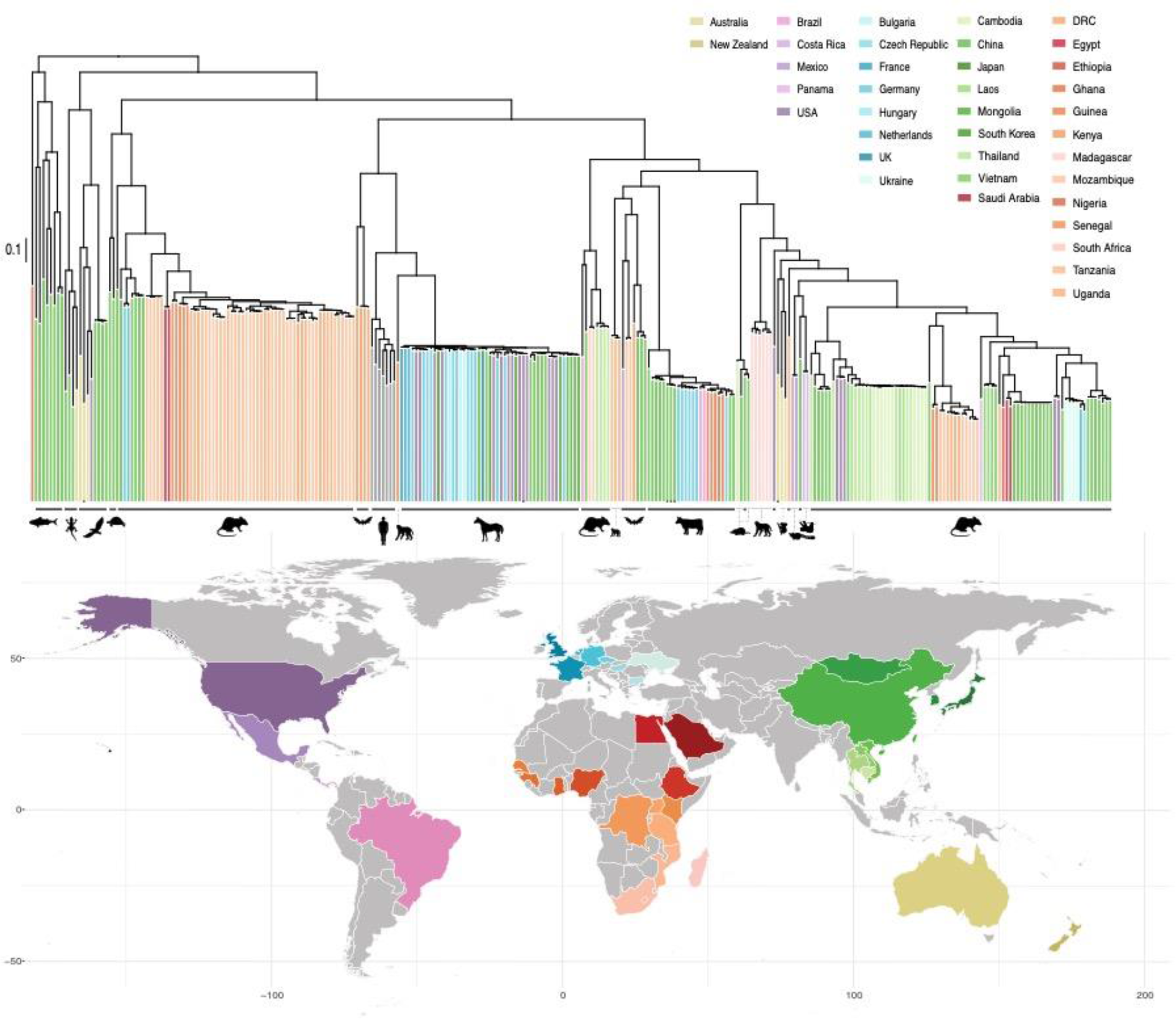
Geographic distribution of hepaciviruses based on complete genome sampling locations. Except for the HCV genotypes, clade colors in the hepacivirus ML tree correspond to geographic locations in the map. Only major host types occupying a clade are marked with icons next to the phylogeny. Arthropods hepaciviruses, which include one tick hepacivirus close to marsupial viruses, three tick hepaciviruses from cattle, one mosquito hepacivirus clustering with bird viruses, as well as one canine hepacivirus within the equine lineage are all marked with black dots at the tips.

To this end, we explored the extent to which hepacivirus diversity is structured by geography and host species using a Bayesian multidimensional scaling (BMDS) approach that employs pairwise host and geographic distances (see Methods, section 2.6). We estimated Pagel’s lambda as a measure of phylogenetic signal for the host and geographic traits, either in a strict Brownian diffusion model or a relaxed random walk model. For the latter, we estimated the standard deviation of the lognormal distribution as a measure of the heterogeneity in the diffusion process in host and geographic space. A 5-fold cross-validation approach was applied to i) all mammalian hepaciviruses in our data set (*n* = 257) and ii) to a subset containing only a single representative for bovine and equine viruses, respectively, as well as for clusters of viruses that were sampled from the same host and country (*n* = 123). This analysis indicated that a one-dimensional model provided the best fit to the data in our BMDS approach, and therefore we consistently adopted this model in all our investigations.

In Table 1, we summarize the posterior estimates for Pagel’s lambda and the lognormal standard deviation of the RRWs for different subsets of the hepacivirus data. These include all non-mammalian hepaciviruses (*n* = 13), all mammalian hepaciviruses (*n* = 257), a mammalian subset with only a single representative for clusters of viruses that were sampled from the same host and country (*n* = 160), a subset of the latter with only a single representative for bovine and equine viruses (*n* = 123), only rodent viruses (*n* = 95) and only rodent viruses excluding those sampled from co-infected individuals (*n* = 68). Pagel’s lambda estimates for mammalian data sets indicate a high phylogenetic signal for both host and geography in our BMDS approach and hence no discriminatory power to distinguish between the two. We obtained substantially different estimates only for the small non-mammalian data set, with higher signal for host structure than for geographical structure. This can be also seen in the colored phylogeny (Fig. 6), where the host trait indeed leads to more variation. For only one mammalian subset, we obtained a somewhat lower phylogenetic signal estimate for geography when using a strict Brownian model. However, when using only a single bovine and equine hepacivirus in the subset, we estimated again a high degree of geographic signal. This indicates a higher degree of hepacivirus spatial dispersal in equids and cattle (Fig. 7), which is likely shaped by human-assisted movement on a global scale.

**Fig. 6.**
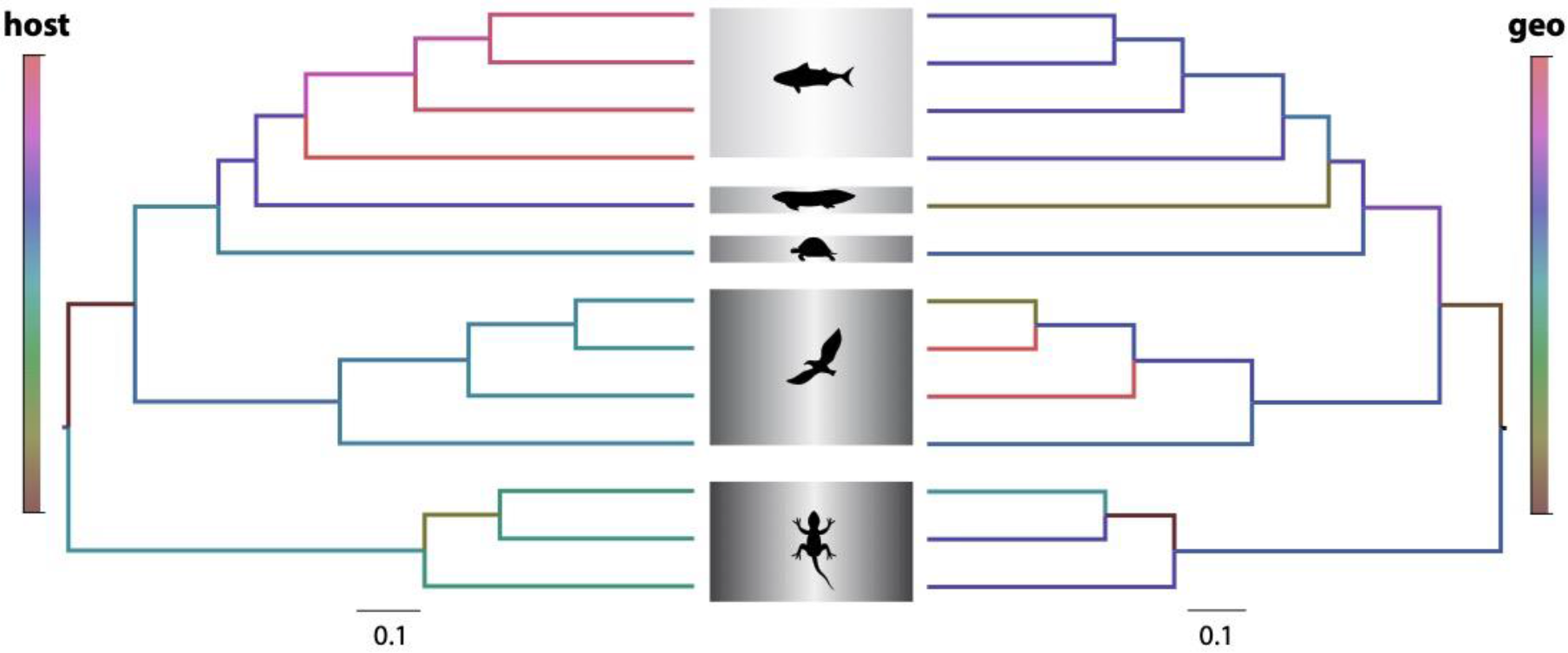
Non-mammalian hepacivirus (*n* = 13) phylogeny colored by precisions of host distances and geographical distances generated from the strict Brownian one-dimensional BMDS analysis. The two phylogenies were derived from the same Bayesian maximum clade credibility (MCC) tree.

**Fig. 7.**
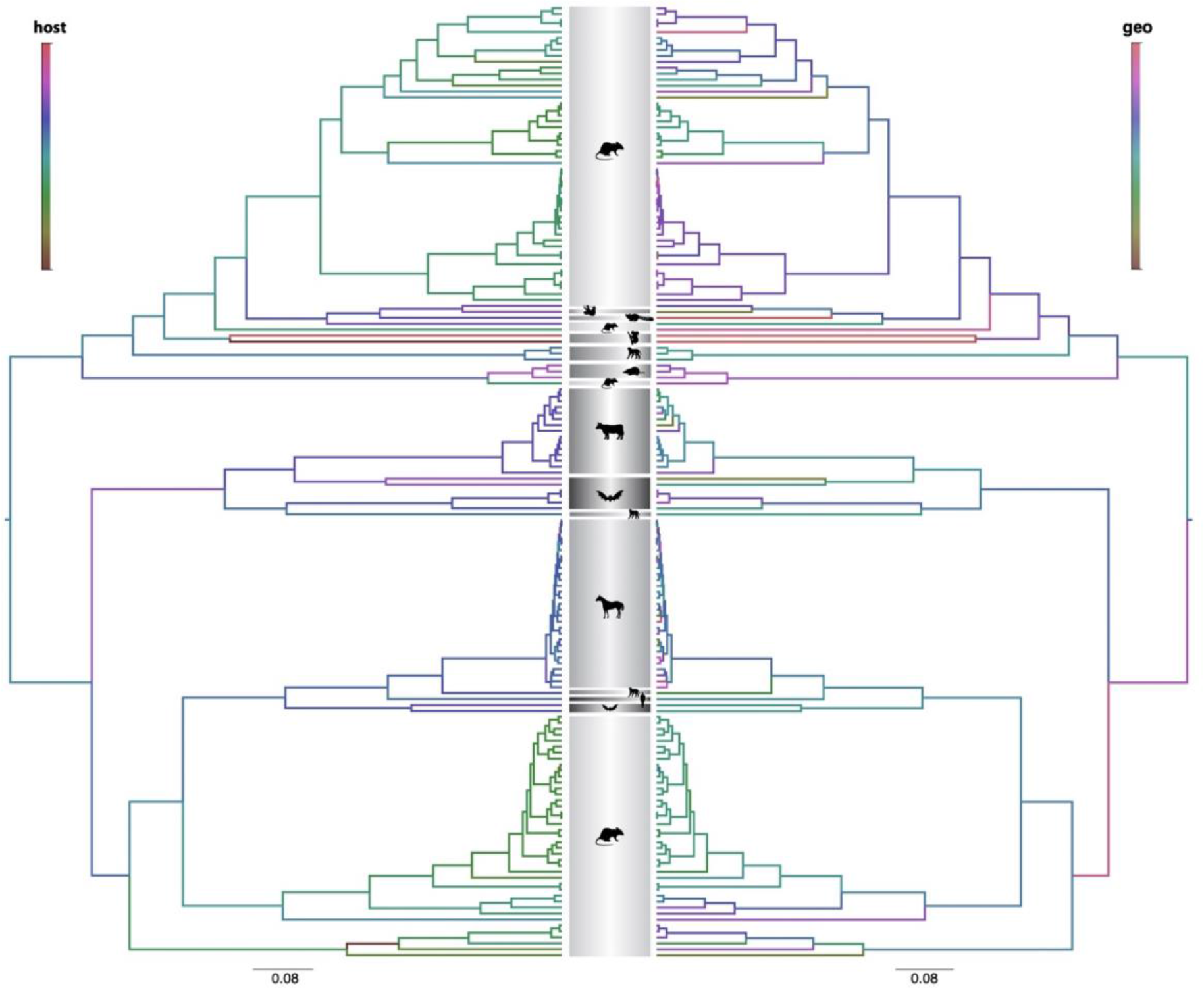
Mammalian hepacivirus (subset 1, *n* = 160) phylogeny colored by precisions of host distances and geographical distances generated from the strict Brownian one-dimensional BMDS analysis. The two phylogenies were derived from the same Bayesian maximum clade credibility (MCC) tree.

**Table 1.** Mean posterior estimates and 95% highest density posterior intervals for Pagel’s lambda and the relaxed random walk standard deviation.

| Data set size |  | Strict Brownian |  | Relaxed random walk |  |  |  |
| --- | --- | --- | --- | --- | --- | --- | --- |
|  |  | Lambda |  | Lambda |  | Standard deviation |  |
|  |  | Host | Geo | Host | Geo | Host | Geo |
| Non-mammalian | 13 | 0.7733<br>(0.3543, 0.9999) | 0.2833<br>(0.0001, 0.7263) | 0.8343<br>(0.4436, 0.9999) | 0.2548<br>(0.0003, 0.684) | 1.9475<br>(0.0263, 4.8118) | 1.7737<br>(0.0076, 4.2552) |
| Mammalian | 257 | 0.9999<br>(0.9996-1.0000) | 0.9999<br>(0.9998-1.0000) | 0.9967<br>(0.9945-0.9987) | 0.9998<br>(0.9997-1.0000) | 1.5855<br>(0.2109, 3.7811) | 23.395<br>(15.8077, 30.5774) |
| Mammalian subset 1* | 160 | 0.9997<br>(0.9991-1.0000) | 0.7383<br>(0.5573-0.8809) | 0.9998<br>(0.9996-1.0000) | 0.9999<br>(0.9976-0.9999) | 5.764<br>(2.8956, 8.6949) | 14.905<br>(8.5009, 19.6000) |
| Mammalian subset 2** | 123 | 0.9996<br>(0.9972-1.0000) | 0.9997<br>(0.9942-1.0000) | 0.9998<br>(0.9993-1.0000) | 0.9999<br>(0.9996-1.0000) | 5.4163<br>(2.7742, 8.5257) | 8.849<br>(5.0008, 13.2999) |
| Rodent | 95 | 0.9996<br>(0.9989-1.0000) | 0.9997<br>(0.9990-1.0000) | 1.0000<br>(0.9999, 1.0000) | 1.0000<br>(0.9999, 1.0000) | 2.1285<br>(0.5031, 4.1033) | 4.041<br>(1.8113, 7.0174) |
| Rodent, no co-infections | 68 | 0.9994<br>(0.9979-1.0000) | 0.9994<br>(0.9980-1.0000) | 0.9998<br>(0.9993-1.0000) | 0.9945<br>(0.9811, 1.0000) | 2.6692<br>(0.8834, 4.9230) | 5.982<br>(3.0039, 10.0134) |

\* Mammalian subset with only single representatives for clusters of viruses that were sampled from the same host and country
\*\* Same as above with only single representatives for the bovine and equine viruses

In contrast to the phylogenetic signal estimates, the RRW standard deviation estimates are markedly different for the diffusion process in host and geographic space in the mammalian virus data sets. Specifically, we obtained consistently higher estimates for the geographic diffusion process indicating more variability or heterogeneity than for the host diffusion process. In the rodent data sets, we estimated less heterogeneity in the geographic diffusion process as compared to the collection of mammalian viruses, but also in these cases the host diffusion process is more regular.

### Divergence dating reveals the deep hepacivirus evolutionary history

To assess our ability to estimate short-term evolutionary rates for hepaciviruses, we selected five lineages from our hepacivirus data set, which included the bovine and equine-canine hepacivirus clades, and three RHV sub-clades with relatively low genetic diversity (Supplementary Fig. 1). In addition, we included genomic data sets for HCV genotypes 1a, 1b and 3a. Prior to testing for temporal signal using the Bayesian Evaluation of Temporal Signal (BETS) approach, we explored the impact of recombination in those datasets. Significant evidence for recombination was detected in the bovine, equine, rodent 1 and rodent 3 hepacivirus lineages (Supplementary Table 8). Recombinant regions were masked and the datasets were further reduced by only keeping sequences with known sampling time.

For each host-specific lineage, we estimated the (log) Bayes factor support for a model that employs sampling time (dated tips) against a model that treats sequences as contemporaneous (same sampling time) This BETS analysis revealed that the equine hepacivirus lineage, as well as HCV genotypes 1a, 1b and 3a exhibit strong support for the presence of temporal signal (log Bayes factor > 3). For the bovine and the three rodent hepacivirus lineages tested, BETS reported evidence against significant temporal signal (Table 2.).

**Table 2.**
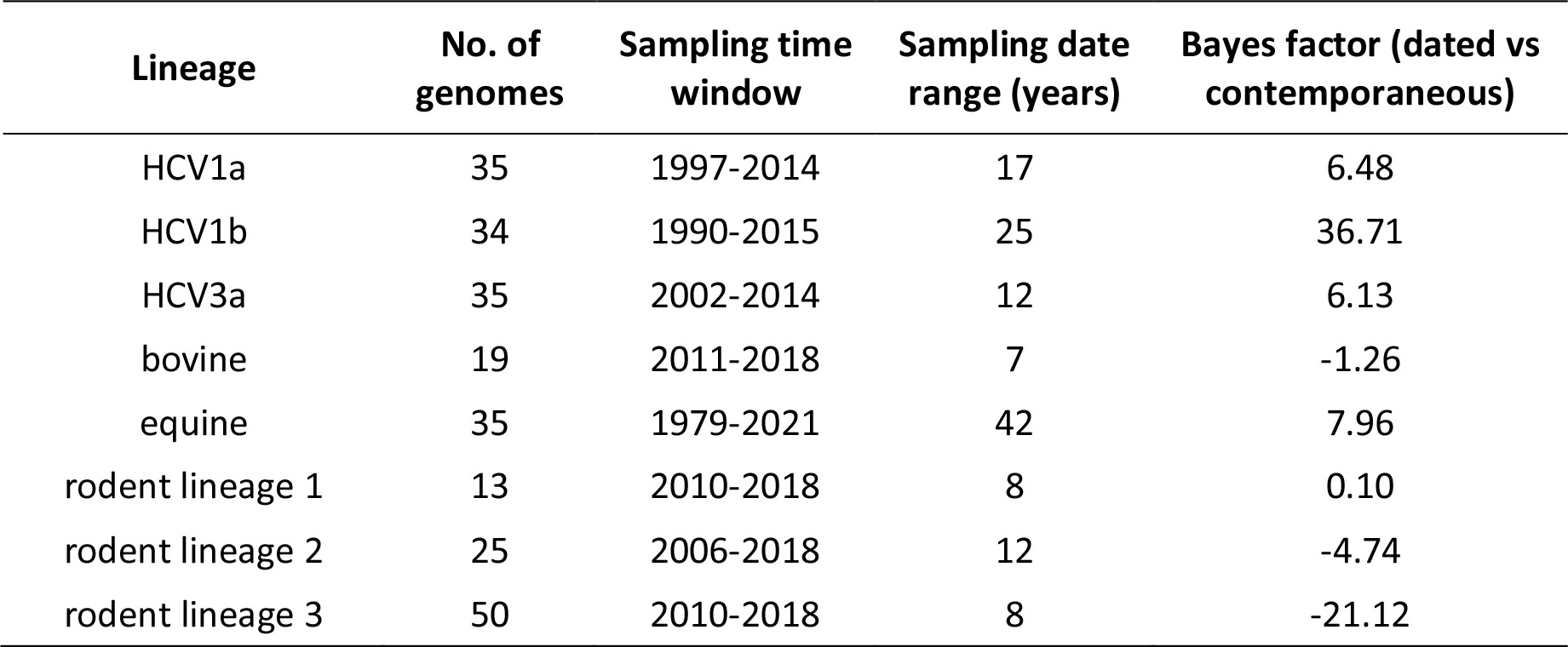
Temporal signal results from BETS.

Using the short-term substitution rate of 4.6 (95% HPD: 3.9 – 5.4) × 10^-4^ substitutions per site per year estimated from HCV genotypes 1a, 1b and 3a, we applied the recently developed mechanistic time-dependent rate decay PoW model to date the hepacivirus evolutionary history (Ghafari et al. 2021). Based on the PoW model estimates, we estimate the tMRCA of the mammalian and non-mammalian hepaciviruses to be 21.9 (17.8 – 27.4) million years ago (MYA), thus, indicating that the origins of the *Hepacivirus* genus occurred at least around that time (Fig. 8 and Table 3). Hepaciviruses from fish, reptiles and avian hosts diverged 19.2 (15.4 – 24.8) MYA, while the mammalian hepaciviruses split dates back to 17.4 (13.8 – 21.8) MYA. Interestingly, this split coincides with the rodent and bat hepacivirus diversification, which concords with a deep evolutionary history of these lineages, driven to some extent by host switching events to other mammalian species.

**Fig. 8.**
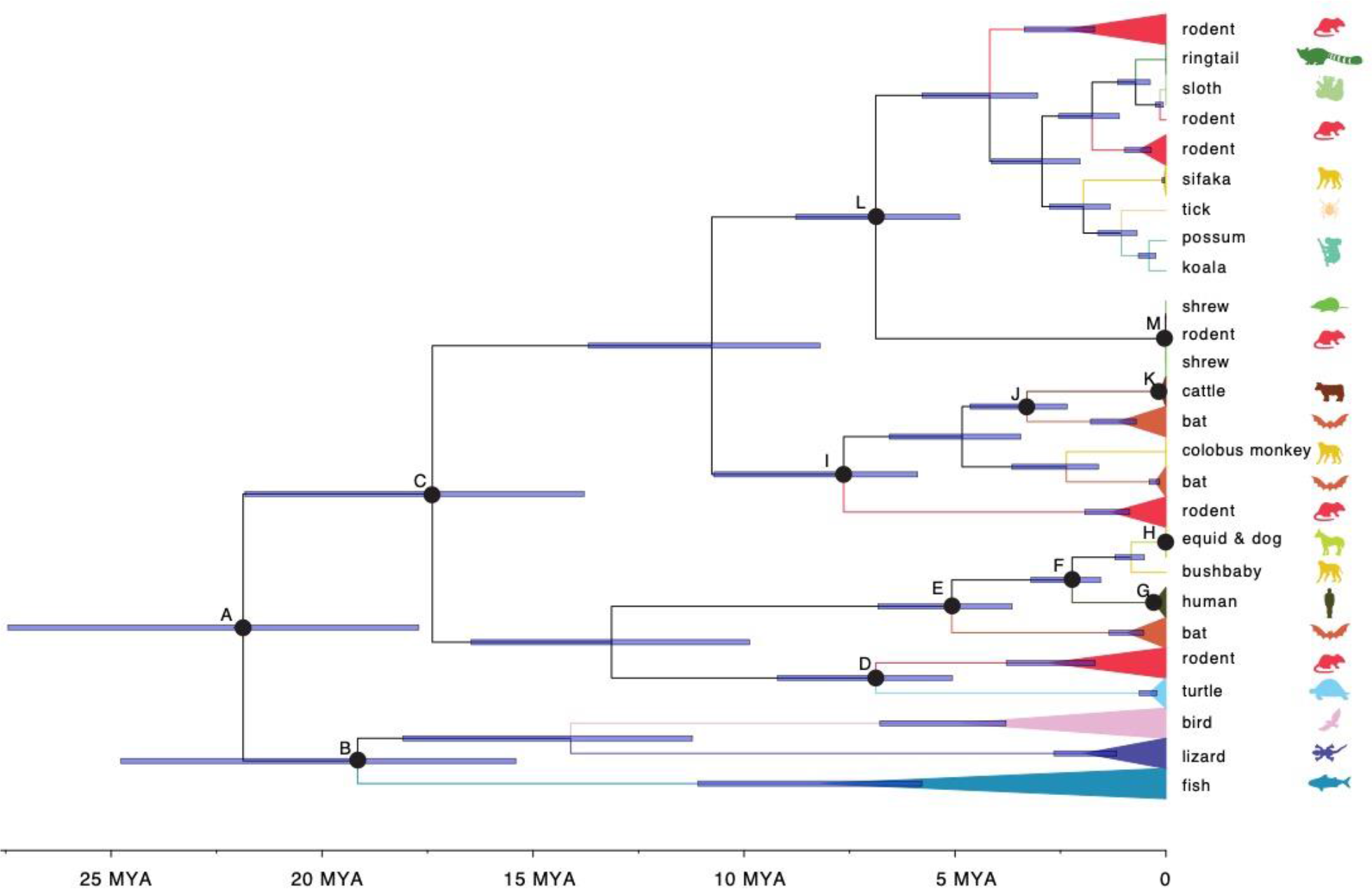
Time-scaled hepaciviruses phylogeny obtained using the PoW-model. Lineages were collapsed based on the host type. Node age uncertainty is shown with 95% highest posterior density (HPD) as interval blue bars. Nodes of interest are marked as A - M for further discussion, and the tMRCA and 95% HPD are listed in Table 3.

**Table. 3.**
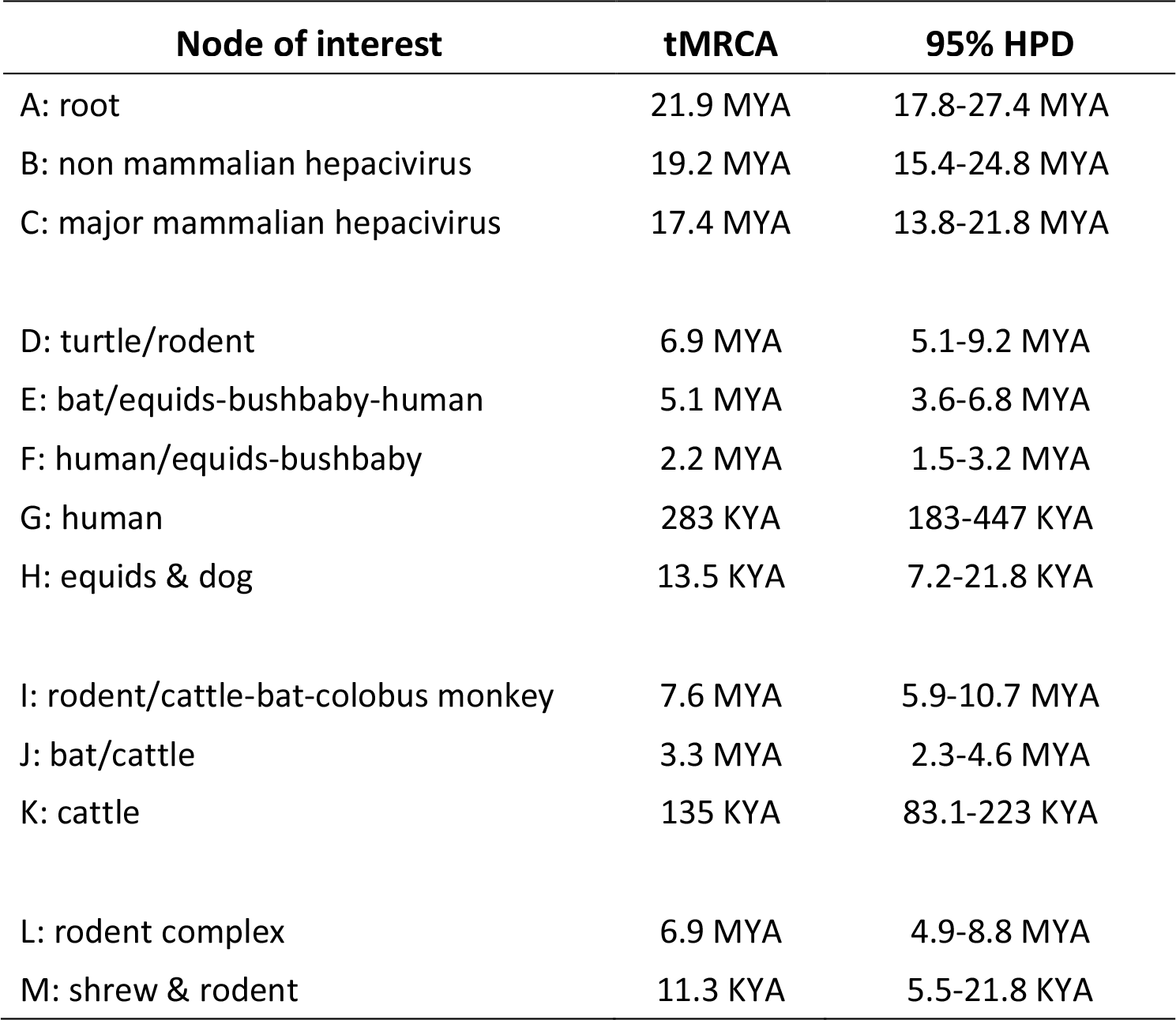
Hepaciviruses divergence times for particular nodes of interest.

When focusing on the descendant clades, bovine, bat and non-human primate hepaciviruses separated from RHVs (node I in Fig. 8) around 7.6 (5.9 – 10.7) MYA, whereas the tMRCA of the diverse mammalian cluster 1 (Fig. 1 – or node L in Fig. 8) is estimated at 6.9 (4.9 – 8.8) MYA. Furthermore, we estimate that the common ancestor of the HCV genotypes dates back to 283 (183 – 447) KYA, in broad agreement with the HCV divergence time estimates reported by Ghafari et al. (2021). Finally, the most recent radiations correspond to the bovine, equine-canine and shrew hepacivirus lineages, which appear to have diversified 135 (83.1 – 223) KYA, 13.5 (7.2 – 21.8) KYA and 11.3 (5.5 – 21.8) KYA, respectively. These time estimates indicate that HCV originated much earlier than the emergence of these hepacivirus lineages, unless HCV was introduced through multiple zoonotic jumps more recently from a hitherto unsampled reservoir.

## Discussion

Animal hepaciviruses have received considerable attention during the last decade because of the search for the zoonotic origin of HCV and the interest in developing surrogate animal models for HCV clinical and vaccine studies (Hartlage et al. 2016; Trivedi et al. 2018). In this work, we screened a comprehensive collection of wild mammalian specimens (*n* = 1,672) mainly from Africa and Asia. Our screening complements previous efforts and extends the host range of hepaciviruses in the rodent population by identifying 13 novel host species and 3 novel host genera, particularly in the Cricetidae and Muridae families. We also contribute to knowledge of the geographic distribution of rodent hepacivirus genomes, not only by extending the sampling locations within China and Africa, but also by including specimens from previously unrepresented locations in the Middle East and Western Africa.

In our screening, we observed that the percentage of hepacivirus positive samples varied considerably among host species. Although this might be biased due to the uneven number of specimens tested per host species, it may also suggest a potentially uneven hepacivirus prevalence in natural rodent populations. The sampling bias and variation of hepacivirus prevalence among different host species should be considered when assessing the spatial distribution of hepaciviruses, especially for locations with high host species diversity.

Our genome-wide virus phylogeny demonstrates that small mammals, especially bats and rodents, constitute an important source of divergent hepaciviruses. The clustering of these virus genomes does not follow the mammalian taxonomy, which hints to a potentially large number of cross-species transmission events during the hepacivirus evolutionary history. Those multiple host-switching events are further supported by the results of our co-phylogenetic analysis, where we observe a high degree of incongruence between virus and host trees, especially between taxa corresponding to shallow nodes. This is in line with previous studies, which estimated a frequency of 65% of all evolutionary events in the hepacivirus and pegivirus genera to represent cross-species transmission (Mifsud et al. 2023). Despite frequent host jumps, the deeper hepacivirus topology appears to reflect an important role of virus-host co-divergence that is also supported by our observation of an overall significant signal of co-divergence. Mifsud et al. (2023) have detected and quantified an accompanying signal of virus-host co-divergence in their analyses (frequency of 22% of all evolutionary events), indicating that hepaciviruses may to some extent have co-diverged with their hosts over longer evolutionary timescales. Taken together, our results corroborate the hypothesis that hepaciviruses have crossed the species barriers multiple times in the past, either in cases of spillover between species from the same genus (Walter et al. 2017; Bletsa et al. 2021) or across different orders (Pybus and Thézé 2016; Moreira-Soto et al. 2020). Relatively frequent cross-species transmission is commonly observed in rapidly-evolving RNA viruses (Geoghegan et al. 2017; Shi et al. 2018).

Due to the wide host spectrum of hepaciviruses and their high genetic heterogeneity, virus phylogenetic relationships have previously been explored mainly from the perspective of their hosts’ diversity (Pybus and Thézé 2016; Porter et al. 2020; Bletsa et al. 2021; Mifsud et al. 2023). However, the spatial component of prevalence can be also crucial in shaping virus diversity. Hepacivirus spatial structure has so far been investigated only at an intraspecies level, for instance, within HCV genotypes (Markov et al. 2009; Iles et al. 2014; Hostager et al. 2019). To compare the extent to which geography and hosts have influenced viral diversification within the *Hepacivirus* genus, we adopted a BMDS approach based on host genetic and geographic distances. This approach has previously been used to assess the antigenic evolution of influenza strains (Bedford et al. 2014; Langat et al. 2017) and to evaluate how mobility processes have shaped viral dispersal (Holbrook et al. 2021). Using a measure of phylogenetic signal did not allow us to meaningfully distinguish between the degree of host and spatial structuring in the mammalian hepaciviruses; both generally exhibited high phylogenetic signal, although some impact of hepaciviruses in livestock was noted that can be explained by human transportation. We note that a phylogenetic signal measure (Pagel’s lambda) close to 1 does not necessarily imply an absolute clustering according to this trait, but that the trait distribution follows the pattern that is expected under a Brownian diffusion process over the phylogeny. As such a high phylogenetic signal measure for the host trait does not necessarily imply strong co-divergence, and it could be compatible with a process of preferential host switching (Charleston and Robertson 2002). Based on a measure of diffusion rate variability in relaxed random walk models, we recover more pronounced differences between host and geographic diffusion processes in mammalian hosts. Specifically, we detect a considerably higher degree of variability in the geographic diffusion process, but it remains challenging to provide a clear interpretation of this.

Contrary to the mammalian hepaciviruses, non-mammalian hepaciviruses show a markedly different pattern with a substantially higher phylogenetic signal according to host compared to geography. The non-mammalian hosts include relatively closely related host types, such as birds or fish, which are globally distributed and thus the host structure component is expected to have a stronger effect on the diffusion process. Moreover, the comparably higher host phylogenetic signal might also be caused by a more prominent virus-host co-divergence process (relative to cross-species transmissions) in the more sparsely sampled non-mammalian hepacivirus evolutionary history (Porter et al. 2020; Mifsud et al. 2023). It is indeed important to acknowledge that the relatively sparse sampling, both in terms of the number of available virus genomes and the variation in sampling locations, of the non-mammalian hepacivirus dataset may impact the host-geography comparative analyses.

Complementing previous results by Bletsa et al. (2021), we find that *Lophuromys* rodents (brush-furred mice) from central and western Africa frequently harbor hepacivirus co-infections. The distribution of hepacivirus co-infection strains within Africa (West, Central and East Africa) seems to be associated with the distribution of the *Lophuromys* rodents and suggests that this finding is most likely restricted to members of this particular rodent genus, although the ecological reason for this is still unclear (Bletsa et al. 2021). The majority of our hepacivirus genomes obtained from co-infected individuals form sister lineages in the phylogenetic tree and provide significant links in the reconciliation analysis. Given the close phylogenetic distance between both the *Lophuromys* hosts and the co-infection strains, it seems reasonable that we have not yet observed any impact caused by the RHV co-infections on either our overall virus-host co-phylogenetic or BMDS analyses, since we obtained similar results with or without including all the RHV genomes from the co-infected individuals.

Using a substitution rate decay model to account for time dependent rates, we formally infer for the first time the timescale for the long-term evolutionary history of the *Hepacivirus* genus from present-day virus genome data only. We estimate that the origin of hepacivirus dates back to about 22 million years ago. In very broad terms, the ancient origin of this virus genus is also implied by previous studies, which suggested an origin of the whole *Flaviviridae* family close to the emergence of metazoans 750-800 million years ago (Bamford et al. 2022; Mifsud et al. 2023). However, those previous estimates include all flavivirid groups (*Flavi-*, *Hepaci-*, *Pegi-* and *Pestivirus* genera) and provide estimates of the evolutionary origins based on virus-host co-divergence hypotheses combined with EVEs calibrations. While our hepacivirus molecular dating estimates cannot be directly compared to the estimates of the whole *Flaviviridae* family, the divergence dates for subclades seems to be in very good agreement with the divergence date of the HCV genotypes (Ghafari et al. 2021).

In terms of other hepacivirus subclades, the presence of mammalian hepaciviruses (17.4 MYA) is estimated to have originated much earlier than the origin of modern humans (0.3 – 1 MYA) (Bergström et al. 2021), thus emphasizing that the virus was possibly circulating in non-human mammalian hosts for millions of years before it was introduced into the human population through a single or multiple cross-species transmission events. The HCV zoonotic source has been often proposed to lie in rodents or bats due to the great genetic hepacivirus diversity circulating in these animals and their general importance as pathogen reservoirs (Drexler et al. 2013; Kapoor et al. 2013; de Souza et al. 2019; Bletsa et al. 2021). Our time tree also demonstrates that rodent hepaciviruses diverged earlier than hepaciviruses from other mammalian hosts and that their divergence overlaps with the MRCA of the entire mammalian hepacivirus group. This suggests a possible rodent origin for many other mammalian hepaciviruses, although the transmission routes from those small mammals to other mammalian hosts remain unknown. Finally, our results do not support the current bovine, equine, canine, or shrew hepaciviruses as the origin of HCV, since HCV is more diversified and diverged earlier than the equine, canine and shrew hepaciviruses. We expect that a more complete picture of the hepacivirus diversity and potential zoonotic reservoirs for HCV may emerge from additional screening efforts. The origin of equine and canine hepacivirus is estimated to be 13.5 KYA, well aligned with the start of agriculture communities development and livestock domesticating (12 - 14 KYA) (Hartung 2013). The bovine hepacivirus originated at 135 KYA, which seems to suggest an hepacivirus introduction into the aurochs population before domesticating. However, the complex domestication patterns and introgression in bovine evolutionary history (Murray et al. 2010; Upadhyay et al. 2017) make the hepacivirus emergence event difficult to interpret. Finally, we acknowledge that our estimates of the time scale are primarily derived from the application of the PoW model on present-day virus sequence data and currently rely on the modern rate of HCV and the maximum substitution rate of ssRNA viruses in general. Given the very deep evolutionary history of hepaciviruses and the overall scarcity of archival molecular data, which are considered genomic fossil records, our results may be sensitive to the type of model applied and they may change when additional data will become available in the future.

In summary, we show that the current phylogeny for members of the *Hepacivirus* genus is largely characterized by a complex combination of micro- and macroevolutionary processes having occurred over timescales of millions of years. We envisage that the methods applied in this study and the novel results obtained will inspire future research on virus evolutionary dynamics and phylogeography of hepaciviruses and, hopefully, lead to uncovering an even wider diversity of circulating animal hepaciviruses.

## Acknowledgement

PL and MAS acknowledge support from the European Research Council (grant agreement no. 725422 – ReservoirDOCS), the National Institutes of Health (R01 AI153044), the Wellcome Trust ARTIC (Collaborators Award 206298/Z/17/Z ARTIC network) & supplement, the Research Foundation - Flanders (Fonds voor Wetenschappelijk Onderzoek - Vlaanderen, G066215N, G0D5117N and G0B9317N) and by HORIZON 2020 EU grant 874850 MOOD. YL acknowledges the support from China Scholarship Council (CSC). MG acknowledges support from the Biotechnology and Biological Science Research Council (BBSRC) grant number BB/M011224/1. AJH acknowledges support from the National Science Foundation (DMS 2236854 and DMS 2152774) and the National Institutes of Health (K25 AI153816). SC and JPW acknowledge support from the Zoonoses and Emerging Livestock Systems programme (grant no. BB/L018985). JGB acknowledges support from the Czech Science Foundation (22-32394S).

## Data availability

All sequences generated in this study have been submitted in the NCBI GenBank database under the following accession numbers:

– 34 rodent hepacivirus genomes (OM161982 - OM161997, OM161999 - OM162013, OM162015 - OM162017)
– 22 rodent cytochrome b sequences (OM324311 - OM324328, OM324330 - OM324333).

## References

Abascal F, Zardoya R, Telford MJ. 2010. TranslatorX: multiple alignment of nucleotide sequences guided by amino acid translations. Nucleic Acids Res 38:W7–W13.

Aiewsakun P, Katzourakis A. 2016. Time-Dependent Rate Phenomenon in Viruses. Ross SR, editor. J Virol 90:7184–7195.

Andrews S. 2010. Babraham Bioinformatics - FastQC A Quality Control tool for High Throughput Sequence Data. Available from: https://www.bioinformatics.babraham.ac.uk/projects/fastqc/

Arenas M, Posada D. 2010. The Effect of Recombination on the Reconstruction of Ancestral Sequences. Genetics 184:1133–1139.

Ayres DL, Darling A, Zwickl DJ, Beerli P, Holder MT, Lewis PO, Huelsenbeck JP, Ronquist F, Swofford DL, Cummings MP, et al. 2012. BEAGLE: An Application Programming Interface and High-Performance Computing Library for Statistical Phylogenetics. Systematic Biology 61:170–173.

Baele G, Lemey P, Suchard MA. 2016. Genealogical Working Distributions for Bayesian Model Testing with Phylogenetic Uncertainty. Syst Biol 65:250–264.

Balbuena JA, Míguez-Lozano R, Blasco-Costa I. 2013. PACo: A Novel Procrustes Application to Cophylogenetic Analysis. PLoS One 8:e61048.

Bamford CGG, de Souza WM, Parry R, Gifford RJ. 2022. Comparative analysis of genome-encoded viral sequences reveals the evolutionary history of flavivirids (family Flaviviridae). Virus Evolution:veac085.

Bankevich A, Nurk S, Antipov D, Gurevich AA, Dvorkin M, Kulikov AS, Lesin VM, Nikolenko SI, Pham S, Prjibelski AD, et al. 2012. SPAdes: A New Genome Assembly Algorithm and Its Applications to Single-Cell Sequencing. J Comput Biol 19:455–477.

Bedford T, Suchard MA, Lemey P, Dudas G, Gregory V, Hay AJ, McCauley JW, Russell CA, Smith DJ, Rambaut A. 2014. Integrating influenza antigenic dynamics with molecular evolution. Losick R, editor. eLife 3:e01914.

Bergström A, Stringer C, Hajdinjak M, Scerri EML, Skoglund P. 2021. Origins of modern human ancestry. Nature 590:229–237.

Bletsa M, Vrancken B, Gryseels S, Boonen I, Fikatas A, Li Y, Laudisoit A, Lequime S, Bryja J, Makundi R, et al. 2021. Molecular detection and genomic characterization of diverse hepaciviruses in African rodents. Virus Evolution 7:veab036.

Bolger AM, Lohse M, Usadel B. 2014. Trimmomatic: a flexible trimmer for Illumina sequence data. Bioinformatics 30:2114–2120.

Bonwitt J, Sáez AM, Lamin Joseph, Ansumana R, Dawson M, Buanie J, Lamin Joyce, Sondufu D, Borchert M, Sahr F, et al. 2017. At Home with Mastomys and Rattus: Human-Rodent Interactions and Potential for Primary Transmission of Lassa Virus in Domestic Spaces. The American Journal of Tropical Medicine and Hygiene 96:935–943.

Breitfeld J, Fischer N, Tsachev I, Marutsov P, Baymakova M, Plhal R, Keuling O, Becher P, Baechlein C. 2022. Expanded Diversity and Host Range of Bovine Hepacivirus— Genomic and Serological Evidence in Domestic and Wild Ruminant Species. Viruses 14:1457.

Bruen TC, Philippe H, Bryant D. 2006. A Simple and Robust Statistical Test for Detecting the Presence of Recombination. Genetics 172:2665–2681.

Buchfink B, Reuter K, Drost H-G. 2021. Sensitive protein alignments at tree-of-life scale using DIAMOND. Nat Methods 18:366–368.

Cagliani R, Forni D, Sironi M. 2019. Mode and tempo of human hepatitis virus evolution. Comput Struct Biotechnol J 17:1384–1395.

Camacho C, Coulouris G, Avagyan V, Ma N, Papadopoulos J, Bealer K, Madden TL. 2009. BLAST+: architecture and applications. BMC Bioinformatics 10:421.

Castiglione S, Serio C, Mondanaro A, Melchionna M, Raia P. 2022. Fast production of large, time-calibrated, informal supertrees with tree.merger. Palaeontology 65:e12588.

Charleston MA, Robertson DL. 2002. Preferential Host Switching by Primate Lentiviruses Can Account for Phylogenetic Similarity with the Primate Phylogeny. Systematic Biology 51:528–535.

Choo Q-L, Kuo G, Weiner AJ, Overby LR, Bradley DW, Houghton M. 1989. Isolation of a cDNA Clone Derived from a Blood-Borne Non-A, Non-B Viral Hepatitis Genome. *Science*, New Series 244:359–362.

Corman VM, Grundhoff A, Baechlein C, Fischer N, Gmyl A, Wollny R, Dei D, Ritz D, Binger T, Adankwah E, et al. 2015. Highly Divergent Hepaciviruses from African Cattle. Ou J-HJ, editor. J Virol 89:5876–5882.

Criscuolo A, Gribaldo S. 2010. BMGE (Block Mapping and Gathering with Entropy): a new software for selection of phylogenetic informative regions from multiple sequence alignments. BMC Evol Biol 10:210.

Danecek P, Bonfield JK, Liddle J, Marshall J, Ohan V, Pollard MO, Whitwham A, Keane T, McCarthy SA, Davies RM, et al. 2021. Twelve years of SAMtools and BCFtools. GigaScience 10:giab008.

Drexler JF, Corman VM, Müller MA, Lukashev AN, Gmyl A, Coutard B, Adam A, Ritz D, Leijten LM, van Riel D, et al. 2013. Evidence for Novel Hepaciviruses in Rodents. Wang D, editor. PLoS Pathog 9:e1003438.

Duchêne S, Holmes EC, Ho SYW. 2014. Analyses of evolutionary dynamics in viruses are hindered by a time-dependent bias in rate estimates. Proc. R. Soc. B. 281:20140732.

Ge X-Y, Li J-L, Yang X-L, Chmura AA, Zhu G, Epstein JH, Mazet JK, Hu B, Zhang W, Peng C, et al. 2013. Isolation and characterization of a bat SARS-like coronavirus that uses the ACE2 receptor. Nature 503:535–538.

Geoghegan JL, Duchêne S, Holmes EC. 2017. Comparative analysis estimates the relative frequencies of co-divergence and cross-species transmission within viral families. Drosten C, editor. PLoS Pathog 13:e1006215.

Ghafari M, Simmonds P, Pybus OG, Katzourakis A. 2021. A mechanistic evolutionary model explains the time-dependent pattern of substitution rates in viruses. Current Biology 31:4689–4696.e5.

Gu Z, Eils R, Schlesner M. 2016. Complex heatmaps reveal patterns and correlations in multidimensional genomic data. Bioinformatics 32:2847–2849.

Hahn BH, Shaw GM, De KM, Cock, Sharp PM. 2000. AIDS as a Zoonosis: Scientific and Public Health Implications. Science 287:607–614.

Hartlage AS, Cullen JM, Kapoor A. 2016. The Strange, Expanding World of Animal Hepaciviruses. Annu. Rev. Virol. 3:53–75.

Hartung J. 2013. A short history of livestock production. In: Aland A, Banhazi T, editors. Livestock housing. The Netherlands: Wageningen Academic Publishers. p. 21–34. Available from: https://www.wageningenacademic.com/doi/10.3920/978-90-8686-771-4_01

Harvey E, Rose K, Eden J-S, Lo N, Abeyasuriya T, Shi M, Doggett SL, Holmes EC. 2019. Extensive Diversity of RNA Viruses in Australian Ticks. Journal of Virology 93:e01358–18.

Holbrook AJ, Lemey P, Baele G, Dellicour S, Brockmann D, Rambaut A, Suchard MA. 2021. Massive parallelization boosts big Bayesian multidimensional scaling. J Comput Graph Stat 30:11–24.

Hostager R, Ragonnet-Cronin M, Murrell B, Hedskog C, Osinusi A, Susser S, Sarrazin C, Svarovskaia E, Wertheim JO. 2019. Hepatitis C virus genotype 1 and 2 recombinant genomes and the phylogeographic history of the 2k/1b lineage. Virus Evolution 5:vez041.

Huang X, Madan A. 1999. CAP3: A DNA Sequence Assembly Program. Genome Res 9:868– 877.

Huson DH, Bryant D. 2006. Application of Phylogenetic Networks in Evolutionary Studies. Molecular Biology and Evolution 23:254–267.

Iles JC, Raghwani J, Harrison GLA, Pepin J, Djoko CF, Tamoufe U, LeBreton M, Schneider BS, Fair JN, Tshala FM, et al. 2014. Phylogeography and epidemic history of hepatitis C virus genotype 4 in Africa. Virology 464–465:233–243.

Irisarri I, Baurain D, Brinkmann H, Delsuc F, Sire J-Y, Kupfer A, Petersen J, Jarek M, Meyer A, Vences M, et al. 2017. Phylotranscriptomic consolidation of the jawed vertebrate timetree. Nat Ecol Evol 1:1370–1378.

Kapoor A, Simmonds P, Gerold G, Qaisar N, Jain K, Henriquez JA, Firth C, Hirschberg DL, Rice CM, Shields S, et al. 2011. Characterization of a canine homolog of hepatitis C virus. Proc. Natl. Acad. Sci. U.S.A. 108:11608–11613.

Kapoor A, Simmonds P, Scheel TKH, Hjelle B, Cullen JM, Burbelo PD, Chauhan LV, Duraisamy R, Sanchez Leon M, Jain K, et al. 2013. Identification of Rodent Homologs of Hepatitis C Virus and Pegiviruses. Moscona A, editor. mBio 4:e00216–13.

Katoh K. 2002. MAFFT: a novel method for rapid multiple sequence alignment based on fast Fourier transform. Nucleic Acids Research 30:3059–3066.

Kesäniemi J, Lavrinienko A, Tukalenko E, Mappes T, Watts PC, Jurvansuu J. 2019. Infection Load and Prevalence of Novel Viruses Identified from the Bank Vole Do Not Associate with Exposure to Environmental Radioactivity. Viruses 12:44.

Langat P, Raghwani J, Dudas G, Bowden TA, Edwards S, Gall A, Bedford T, Rambaut A, Daniels RS, Russell CA, et al. 2017. Genome-wide evolutionary dynamics of influenza B viruses on a global scale. PLoS Pathog 13:e1006749.

Langmead B, Salzberg SL. 2012. Fast gapped-read alignment with Bowtie 2. Nat Methods 9:357–359.

Larsson A. 2014. AliView: a fast and lightweight alignment viewer and editor for large datasets. Bioinformatics 30:3276–3278.

Lemey P, Rambaut A, Welch JJ, Suchard MA. 2010. Phylogeography Takes a Relaxed Random Walk in Continuous Space and Time. Molecular Biology and Evolution 27:1877–1885.

Markov PV, Pepin J, Frost E, Deslandes S, Labbe A-C, Pybus OG. 2009. Phylogeography and molecular epidemiology of hepatitis C virus genotype 2 in Africa. Journal of General Virology 90:2086–2096.

Martin DP, Lemey P, Posada D. 2011. Analysing recombination in nucleotide sequences. Molecular Ecology Resources 11:943–955.

Martin DP, Murrell B, Golden M, Khoosal A, Muhire B. 2015. RDP4: Detection and analysis of recombination patterns in virus genomes. Virus Evolution [Internet] 1. Available from: https://academic.oup.com/ve/ve/article/2568683/RDP4:

Meier-Kolthoff JP, Auch AF, Huson DH, Goker M. 2007. COPYCAT : cophylogenetic analysis tool. Bioinformatics 23:898–900.

Mifsud JCO, Costa VA, Petrone ME, Marzinelli EM, Holmes EC, Harvey E. 2023. Transcriptome mining extends the host range of the *Flaviviridae* to non-bilaterians. Virus Evolution 9:veac124.

Moreira-Soto A, Arroyo-Murillo F, Sander A-L, Rasche A, Corman V, Tegtmeyer B, Steinmann E, Corrales-Aguilar E, Wieseke N, Avey-Arroyo J, et al. 2020. Cross-order host switches of hepatitis C-related viruses illustrated by a novel hepacivirus from sloths. Virus Evolution 6:veaa033.

Murray C, Huerta-Sanchez E, Casey F, Bradley DG. 2010. Cattle demographic history modelled from autosomal sequence variation. Philos Trans R Soc Lond B Biol Sci 365:2531–2539.

Nguyen L-T, Schmidt HA, von Haeseler A, Minh BQ. 2015. IQ-TREE: A Fast and Effective Stochastic Algorithm for Estimating Maximum-Likelihood Phylogenies. Molecular Biology and Evolution 32:268–274.

Olayemi A, Cadar D, Magassouba N, Obadare A, Kourouma F, Oyeyiola A, Fasogbon S, Igbokwe J, Rieger T, Bockholt S, et al. 2016. New Hosts of The Lassa Virus. Sci Rep 6:25280.

Pagel M. 1999. Inferring the historical patterns of biological evolution. Nature 401:877–884.

Porter AF, Pettersson JH-O, Chang W-S, Harvey E, Rose K, Shi M, Eden J-S, Buchmann J, Moritz C, Holmes EC. 2020. Novel hepaci- and pegi-like viruses in native Australian wildlife and non-human primates. Virus Evolution 6:veaa064.

Pybus OG, Thézé J. 2016. Hepacivirus cross-species transmission and the origins of the hepatitis C virus. Current Opinion in Virology 16:1–7.

Quan P-L, Firth C, Conte JM, Williams SH, Zambrana-Torrelio CM, Anthony SJ, Ellison JA, Gilbert AT, Kuzmin IV, Niezgoda M, et al. 2013. Bats are a major natural reservoir for hepaciviruses and pegiviruses. Proc. Natl. Acad. Sci. U.S.A. 110:8194–8199.

Rambaut A, Lam TT, Max Carvalho L, Pybus OG. 2016. Exploring the temporal structure of heterochronous sequences using TempEst (formerly Path-O-Gen). Virus Evol 2:vew007.

Revell LJ. 2012. phytools: an R package for phylogenetic comparative biology (and other things). Methods in Ecology and Evolution 3:217–223.

Santichaivekin S, Yang Q, Liu J, Mawhorter R, Jiang J, Wesley T, Wu Y-C, Libeskind-Hadas R. 2021. eMPRess: a systematic cophylogeny reconciliation tool. Russell S, editor. Bioinformatics 37:2481–2482.

Schierup MH, Hein J. 2000. Consequences of Recombination on Traditional Phylogenetic Analysis. Genetics 156:879–891.

Schmid J, Rasche A, Eibner G, Jeworowski L, Page RA, Corman VM, Drosten C, Sommer S. 2018. Ecological drivers of Hepacivirus infection in a neotropical rodent inhabiting landscapes with various degrees of human environmental change. Oecologia 188:289– 302.

Schmieder R, Edwards R. 2011. Quality control and preprocessing of metagenomic datasets. Bioinformatics 27:863–864.

Shao J-W, Guo L-Y, Yuan Y-X, Ma J, Chen J-M, Liu Q. 2021. A Novel Subtype of Bovine Hepacivirus Identified in Ticks Reveals the Genetic Diversity and Evolution of Bovine Hepacivirus. Viruses 13:2206.

Sharp PM, Hahn BH. 2011. Origins of HIV and the AIDS Pandemic. Cold Spring Harbor Perspectives in Medicine 1:a006841–a006841.

Shi M, Lin X-D, Chen X, Tian J-H, Chen L-J, Li K, Wang W, Eden J-S, Shen J-J, Liu L, et al. 2018. The evolutionary history of vertebrate RNA viruses. Nature 556:197–202.

Smith DB, Becher P, Bukh J, Gould EA, Meyers G, Monath T, Muerhoff AS, Pletnev A, Rico-Hesse R, Stapleton JT, et al. 2016. Proposed update to the taxonomy of the genera Hepacivirus and Pegivirus within the Flaviviridae family. Journal of General Virology 97:2894–2907.

de Souza W, Fumagalli M, Sabino-Santos G, Motta Maia F, Modha S, Teixeira Nunes M, Murcia P, Moraes Figueiredo L. 2019. A Novel Hepacivirus in Wild Rodents from South America. Viruses 11:297.

Stamatakis A, Auch AF, Meier-Kolthoff J, Göker M. 2007. AxPcoords & parallel AxParafit: statistical co-phylogenetic analyses on thousands of taxa. BMC Bioinformatics 8:405.

Suchard MA, Lemey P, Baele G, Ayres DL, Drummond AJ, Rambaut A. 2018. Bayesian phylogenetic and phylodynamic data integration using BEAST 1.10. Virus Evolution 4:vey016.

Tamura K, Stecher G, Kumar S. 2021. MEGA11: Molecular Evolutionary Genetics Analysis Version 11. Mol Biol Evol 38:3022–3027.

Těšíková J, Bryjová A, Bryja J, Lavrenchenko LA, Goüy de Bellocq J. 2017. Hantavirus Strains in East Africa Related to Western African Hantaviruses. Vector-Borne and Zoonotic Diseases 17:278–280.

Tomlinson JE, Kapoor A, Kumar A, Tennant BC, Laverack MA, Beard L, Delph K, Davis E, Schott Ii H, Lascola K, et al. 2019. Viral testing of 18 consecutive cases of equine serum hepatitis: A prospective study (2014-2018). J Vet Intern Med 33:251–257.

Trivedi S, Murthy S, Sharma H, Hartlage AS, Kumar A, Gadi SV, Simmonds P, Chauhan LV, Scheel TKH, Billerbeck E, et al. 2018. Viral persistence, liver disease, and host response in a hepatitis C–like virus rat model. Hepatology 68:435–448.

Upadhyay MR, Chen W, Lenstra JA, Goderie CRJ, MacHugh DE, Park SDE, Magee DA, Matassino D, Ciani F, Megens H-J, et al. 2017. Genetic origin, admixture and population history of aurochs (Bos primigenius) and primitive European cattle. Heredity 118:169–176.

Upham NS, Esselstyn JA, Jetz W. 2019. Inferring the mammal tree: Species-level sets of phylogenies for questions in ecology, evolution, and conservation. Tanentzap AJ, editor. PLoS Biol 17:e3000494.

Van Nguyen D, Van Nguyen C, Bonsall D, Ngo T, Carrique-Mas J, Pham A, Bryant J, Thwaites G, Baker S, Woolhouse M, et al. 2018. Detection and Characterization of Homologues of Human Hepatitis Viruses and Pegiviruses in Rodents and Bats in Vietnam. Viruses 10:102.

Vrancken B, Lemey P, Rambaut A, Bedford T, Longdon B, Günthard HF, Suchard MA. 2015. Simultaneously estimating evolutionary history and repeated traits phylogenetic signal: applications to viral and host phenotypic evolution. Methods in Ecology and Evolution 6:67–82.

Walter S, Rasche A, Moreira-Soto A, Pfaender S, Bletsa M, Corman VM, Aguilar-Setien A, García-Lacy F, Hans A, Todt D, et al. 2017. Differential Infection Patterns and Recent Evolutionary Origins of Equine Hepaciviruses in Donkeys. Ou J-HJ, editor. J Virol 91:e01711–16.

Wickham H. 2009. ggplot2. New York, NY: Springer Available from: http://link.springer.com/10.1007/978-0-387-98141-3

Wu Z, Han Y, Liu B, Li H, Zhu G, Latinne A, Dong J, Sun L, Su H, Liu L, et al. 2021. Decoding the RNA viromes in rodent lungs provides new insight into the origin and evolutionary patterns of rodent-borne pathogens in Mainland Southeast Asia. Microbiome 9:18.

Wu Z, Lu L, Du J, Yang L, Ren X, Liu B, Jiang J, Yang J, Dong J, Sun L, et al. 2018. Comparative analysis of rodent and small mammal viromes to better understand the wildlife origin of emerging infectious diseases. Microbiome 6:178.

Yu G, Smith DK, Zhu H, Guan Y, Lam TT-Y. 2017. ggtree: an r package for visualization and annotation of phylogenetic trees with their covariates and other associated data. Methods in Ecology and Evolution 8:28–36.

Zaharia M, Bolosky WJ, Curtis K, Fox A, Patterson D, Shenker S, Stoica I, Karp RM, Sittler T. 2011. Faster and More Accurate Sequence Alignment with SNAP. Available from: http://arxiv.org/abs/1111.5572

