## Supplementary Fig.1 for "The evolutionary history of hepaciviruses"

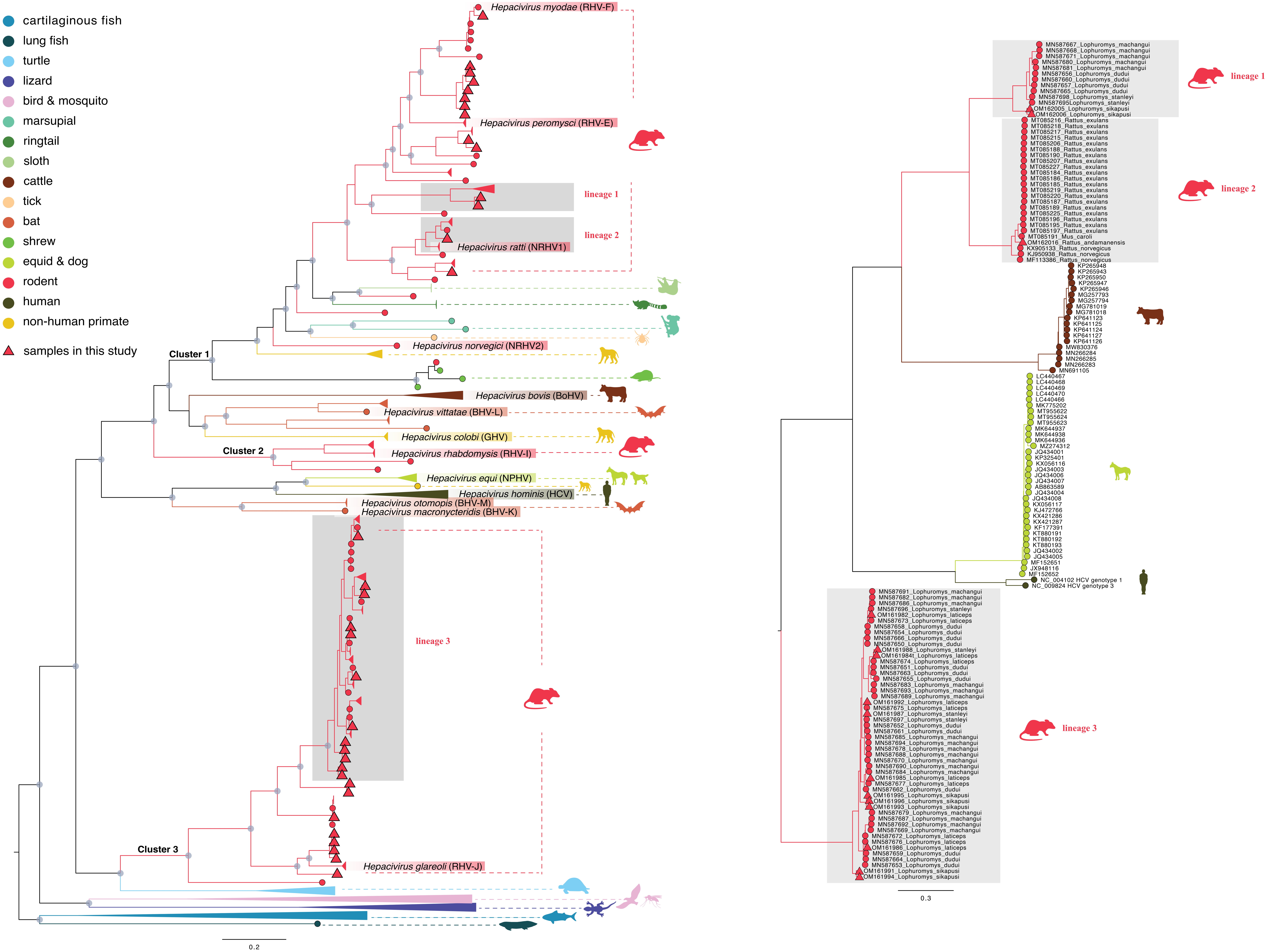


**Supplementary Fig. 1. The hepacivirus lineages that have been selected for assessing recombination and temporal signal according to genetic diversity.**
The subset consists of: HCV genotype 1a (n = 35), HCV genotype 1b (n = 34), HCV genotype 3a (n = 35), cattle (n = 19), equids (n = 35), and 3 lineages for rodent (n = 13, n = 25 and n = 50).
