## Supplementary Fig.2 for "The evolutionary history of hepaciviruses"

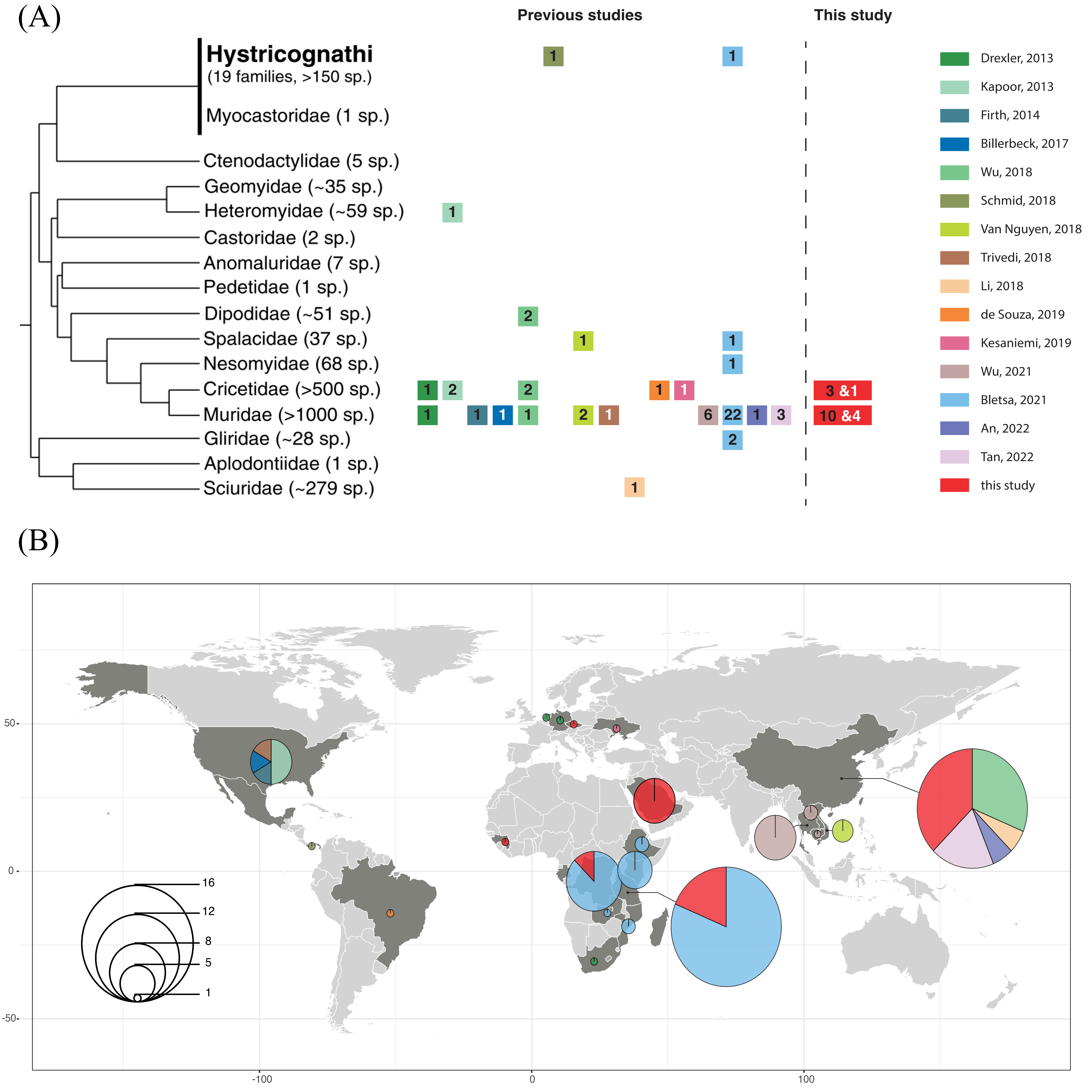


**Supplementary Fig. 2. Overview of hepacivirus screening results in rodents:** (A) The number of hepacivirus-positive host species identified per rodent family to date is summarized in the Rodentia order phylogeny (adapted with permission from Pybus and Thézé 2016). Colors represent the different studies on rodent hepaciviruses. Within the colored boxes, black numbers indicate the number of species that have been newly identified as hepacivirus hosts in the present study, while white numbers indicate the number of species that have already been identified with hepaciviruses in previous studies. (B) Overview of the spatial distribution of locations where rodent hepaciviruses were detected. Countries sampled for hepacivirus are highlighted in dark grey. Pie charts represent the number of hepaci-positive rodent species detected in each locality.
