## Supplementary Fig.3 for "The evolutionary history of hepaciviruses"

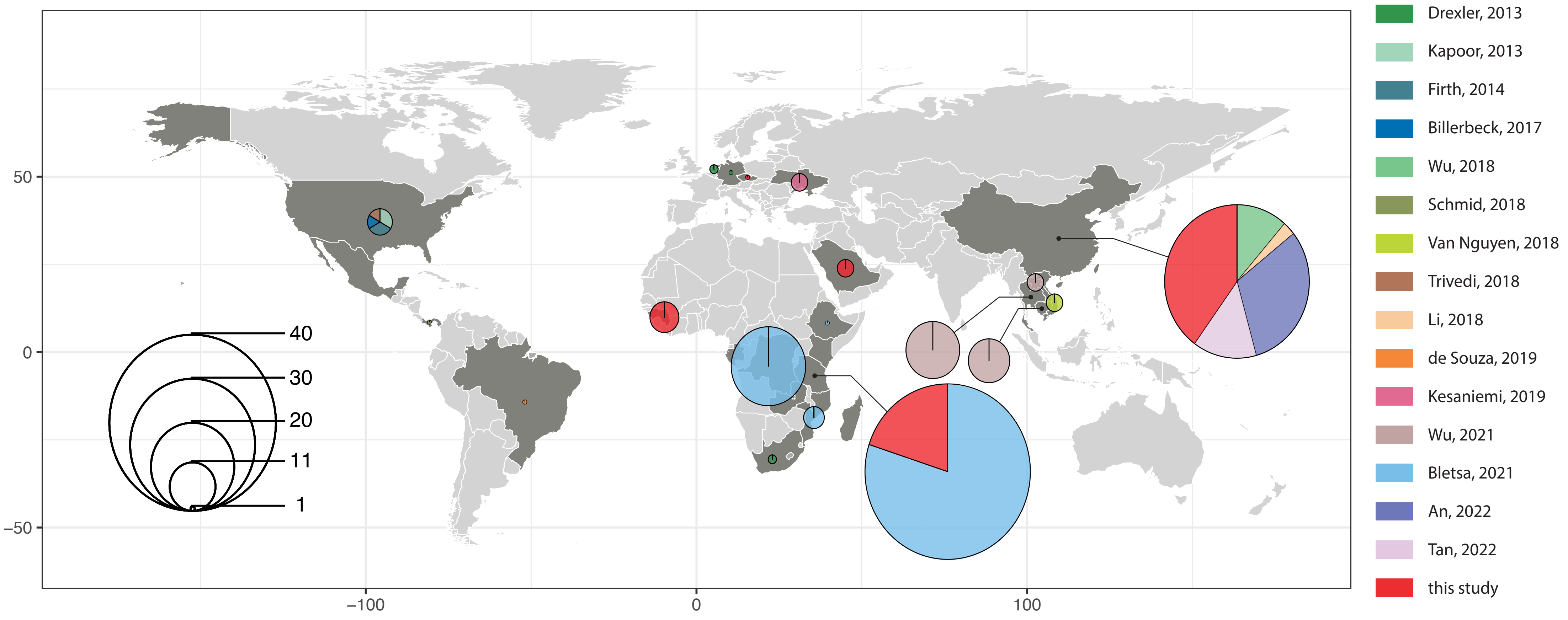


**Supplementary Fig. 3. Overview of the spatial distribution of complete rodent hepacivirus genomes.** Pie charts represent the number of complete hepacivirus genomes generated by this study (red) and previous studies in currently sampled countries.
