## Supplementary Fig.4 for "The evolutionary history of hepaciviruses"

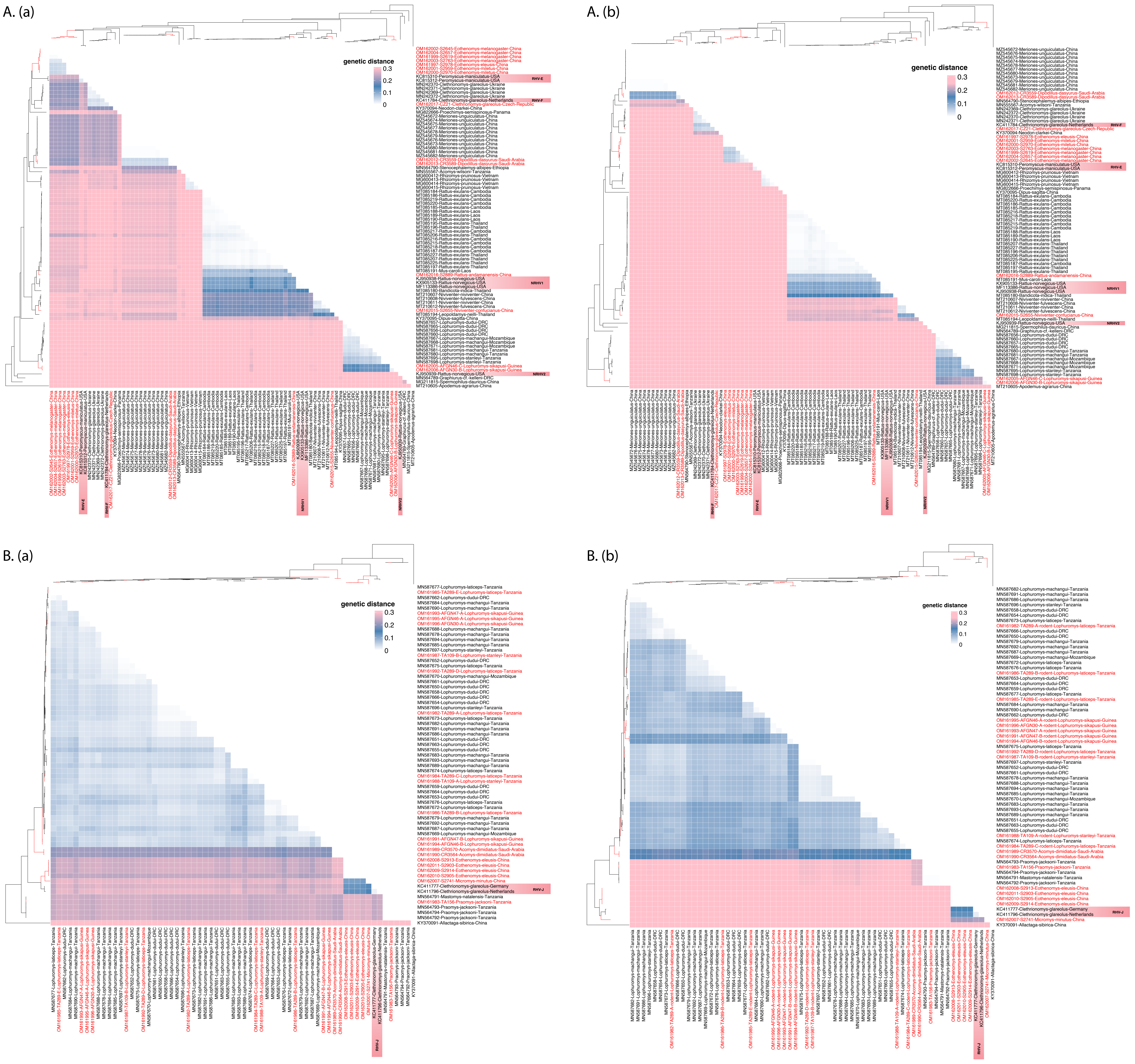


**Supplementary Fig. 2. Amino acid pairwise genetic p-distances heatmap for all rodent hepaciviruses within cluster1 and cluster3 based on NS5B (aa position 2536–2959, NCBI no. AAA45676 as reference) and NS3 (aa position 1123–1566, NCBI no. AAA45676 as reference) region:**
A (a): genetic p-distances heatmap of rodent hepacivirus within cluster 1 on NS5B region, values > 0.3 are marked as pink; A (b): genetic p-distances of rodent hepacivirus within cluster 1 on NS3 region, values > 0.25 are marked as pink;
B (a): genetic p-distances heatmap of rodent hepacivirus within cluster 3 on NS5B region, values > 0.3 are marked as pink; B (b): genetic p-distances of rodent hepacivirus within cluster 3 on NS3 region, values > 0.25 are marked as pink.
