## Supplementary Fig.5 for "The evolutionary history of hepaciviruses"

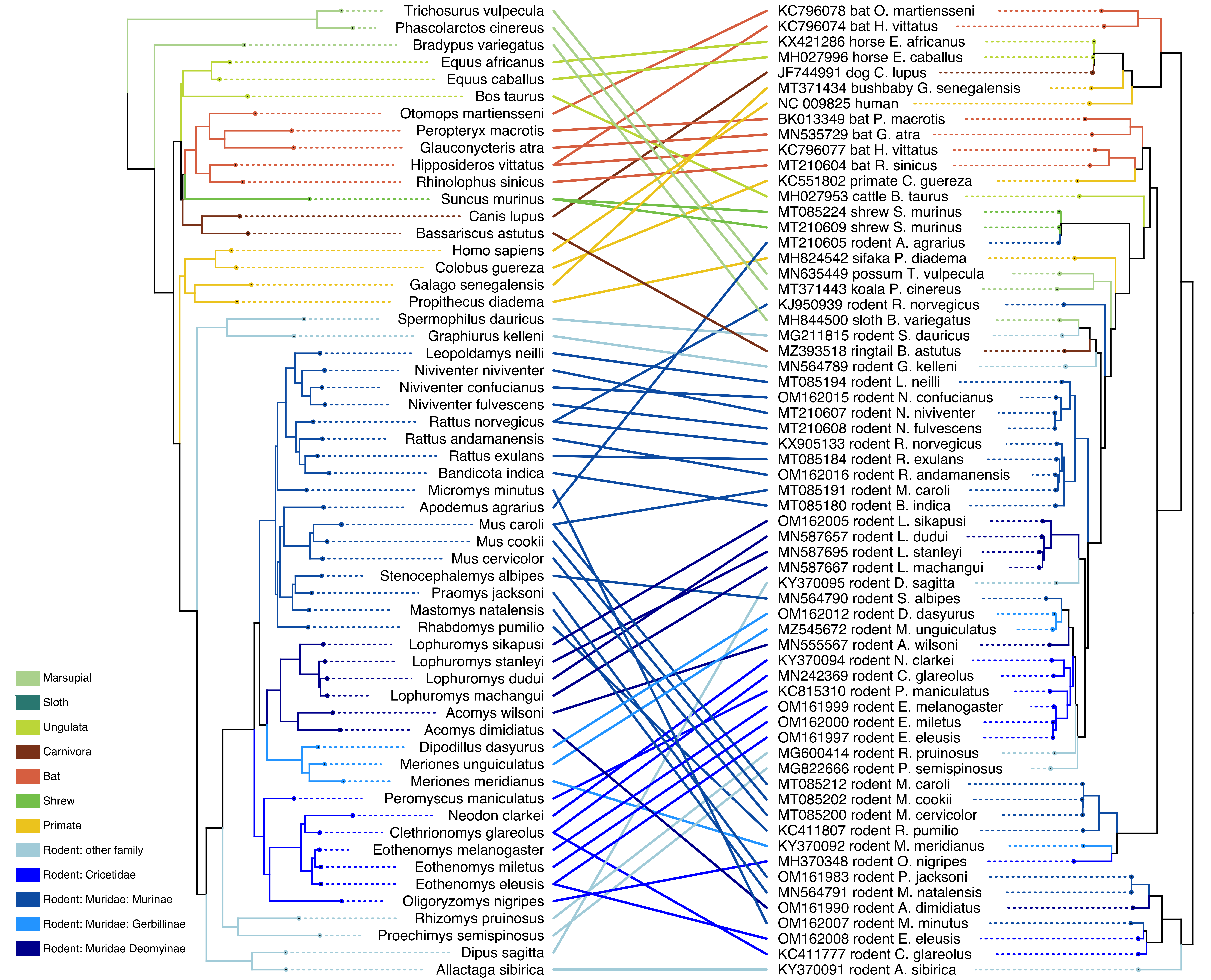


**Supplementary Fig. 3. Tanglegram of host (left) and hepaciviruses (right) after removing co-infections in Lophuromys mice.** The host phylogeny was inferred for 31 genes from 57 mammalian species. For *Myodes glareolus*, *Gerbillus dasyurus* and *Microtus clarkei* we used their updated species names: *Clethrionomys glareolus*, *Dipodillus dasyurus* and *Neodon clarkei*, respectively. The hepacivirus phylogeny was inferred for 63 representative genomes. Clades and associations were colored based on host type.
