## Supplementary Fig.6 for "The evolutionary history of hepaciviruses"

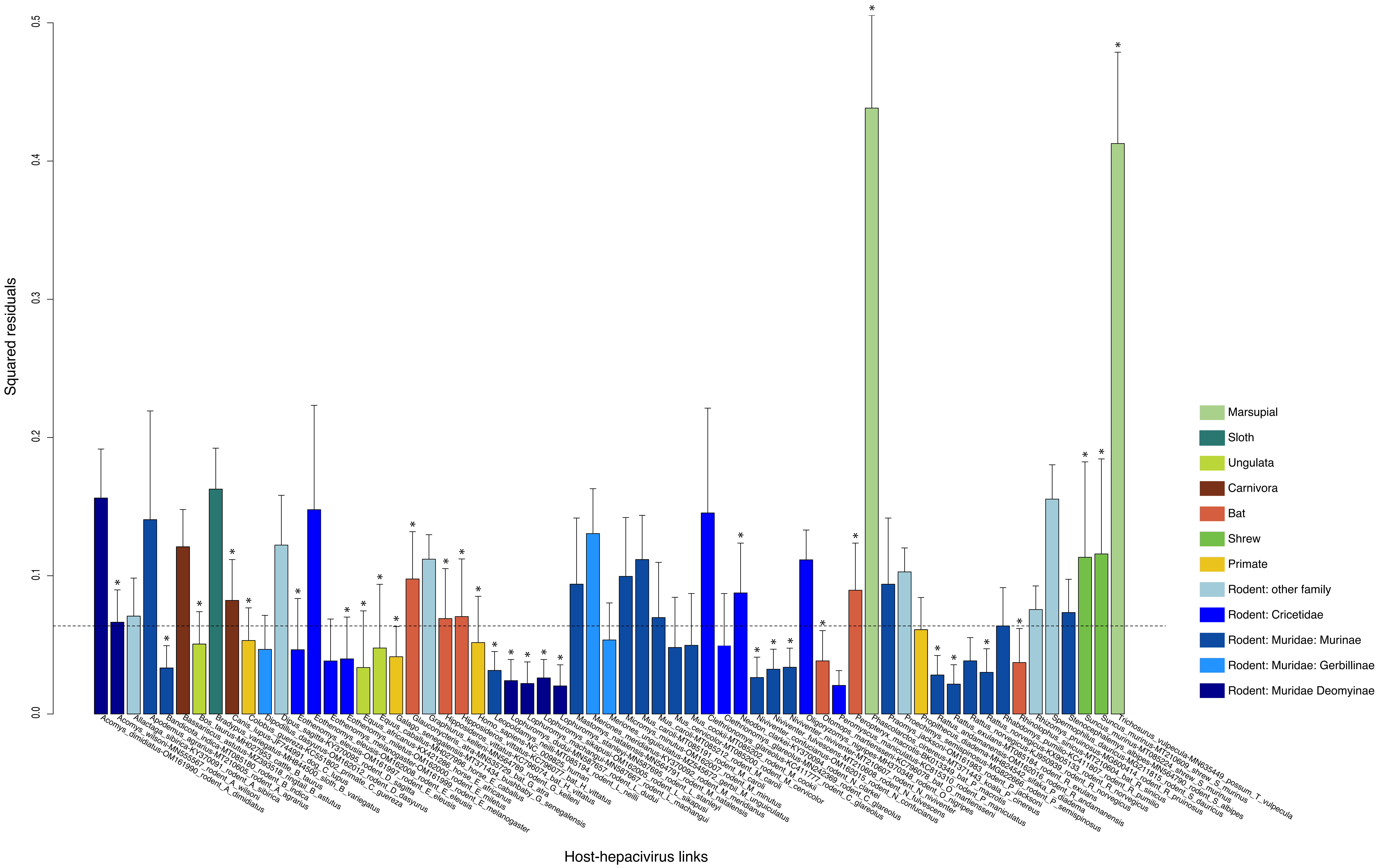


**Supplementary Fig. 4. Contribution of each host-hepacivirus association to the general co-evolution pattern after removing co-infections in Lophuromys mice.**
Bars represent Jack-knifed squared residuals with the upper 95% confidence intervals from PACo test. Stars represent the host-hepacivirus links that were also tested to be significant in AxParafit ($p < 0.05$).
The median squared residual value is shown with a dashed line. Colors represent different host types as shown in Fig. 2.6.
