## Supplementary Table 2 for "The evolutionary history of hepaciviruses"

**Supplementary table S2:** Primer information used in this study.

| **Name** | **Sequence (5'-3')** | **Amplicon size (bp)** | **Study** |
| --- | --- | --- | --- |
| AK4340F1 | GTACTTGCTACTGCNACNCC | 299 | Kapoor et al., 2013 |
| AK4360R1 | TACCCTGTCATAAGGGCRTC |  | Kapoor et al., 2013 |
| AK4340F2 | CTTGCTACTGCNACNCCWCC | 296 | Kapoor et al., 2013 |
| AK4360R2 | TACCCTGTCATAAGGGCRTCNGT |  | Kapoor et al., 2013 |
| qPCR_12SrRNA _512F | AACTCAAAGGACTTGGCGGT | 183 | This study |
| qPCR_12SrRNA _694R | GGCTACACCTTGACCTAACGT |  | This study |
| **Sample specific primers** |  |  |  |
| COVROD3559_2280F | TGTAATTATAGCTTTACTCGCTGC | 351 | This study |
| COVROD3559_2630R | GTTAGCACAAGGAGAGGTACG |  | This study |
| COVROD3564_2302F | TGGGTATCTCCAGTTCAGCT | 311 | This study |
| COVROD3564_2613R | AGTCCTTGCTCTAAATAGAAAGGT |  | This study |
| COVROD3589_527F | TGGCTGTCGGTACTGGTAGA | 493 | This study |
| COVROD3589_1019R | GTCACGTCCGTGCAATATGC |  | This study |
| COVROD3589_904F | GGCACAGTGGGAATTCACCT | 289 | This study |
| COVROD3589_1192R | CCGCCAAGTCACAGACGAAT |  | This study |
| COVROD3589_1860F | GACTGCCTGCTCGTCTATGG | 188 | This study |
| COVROD3589_2047R | TTGACCCCGAATCCCGATTG |  | This study |
| COVROD3589_2123F | CTCCCAGGGTAGCTGGATGT | 434 | This study |
| COVROD3589_2556R | CGAAAGCCACGTCTGACAAG |  | This study |
| COVROD3589_2975F | TACTACCCCTTCGCCACAGA | 469 | This study |
| COVROD3589_3443R | GGAAGAACTCCGTGCAACTG |  | This study |
| COVROD3589_3493F | GTACGTCAACACCAGTCCGT | 434 | This study |
| COVROD3589_3926R | ATGTGAGCCTGGATCCGTTG |  | This study |
| COVROD3589_4092F | CTCCTCCAGGCACTCCAATG | 361 | This study |
| COVROD3589_4452R | AACGTGGGACTCATCGTGAC |  | This study |
| COVROD3589_4630F | CGAAATCGCCACAATGCTGT | 539 | This study |
| COVROD3589_5168R | CAAAGCCCTCCCACTTGCTA |  | This study |
| COVROD3589_5093F | ACTGCAGCAAAGGTGGAACT | 477 | This study |
| COVROD3589_5560R | GAGGCAAGTCGAGGTAGCTG |  | This study |
| COVROD3589_6893F | CGGAAGCGGGTTACCATCAA | 292 | This study |
| COVROD3589_7184R | CCACACGCGCATTGATACTG |  | This study |
| COVROD3589_8039F | GGCTTGTTCGCCTCCTACAT | 453 | This study |
| COVROD3589_8491R | AAGTGCTTGACCCACTCGAG |  | This study |
