## Supplementary Table 3 for "The evolutionary history of hepaciviruses"

**Supplementary table S3:** Reference genomes used to subtract host background reads.

| **NCBI Accession** | **Organism** | **type** | **genus** | **family** |
| --- | --- | --- | --- | --- |
| PVKS01 | Aplodontia rufa | rodent | Aplodontia | Aplodontiidae |
| MTKA01 | Castor canadensis | rodent | Castor | Castoridae |
| AFTD01 | Cricetulus griseus | rodent | Cricetulus | Cricetidae |
| PVKW01 | Orientallactaga bullata | rodent | Orientallactaga | Dipodidae |
| PVJB01 | Muscardinus avellanarius | rodent | Muscardinus | Gliridae |
| ABRO02 | Dipodomys ordii | rodent | Dipodomys | Heteromyidae |
| AEKQ02 | Mus musculus | rodent | Mus | Muridae |
| PVKD01 | Cricetomys gambianus | rodent | Cricetomys | Nesomyidae |
| AABR07 | Rattus norvegicus | rodent | Rattus | Muridae |
| PTPW01 | Ictidomys tridecemlineatus | rodent | Ictidomys | Sciuridae |
| AXCS01 | Nannospalax galili | rodent | Nannospalax | Spalacidae |
| GRCh38.p13 | Homo sapiens | human | Homo | Hominidae |
