## Supplementary Table 7 for "The evolutionary history of hepaciviruses"

**Supplementary table S7:** Hepacivirus positive samples recorded in this study.

| **Name** | **Species** | **Genus** | **Family** | **Country** | **Sample Type** |
| --- | --- | --- | --- | --- | --- |
| S2903 | Eothenomys eleusis | Eothenomys | Cricetidae | China, Asia | tissue (liver) |
| S2913 | Eothenomys eleusis | Eothenomys | Cricetidae | China, Asia | tissue (liver) |
| S2978 | Eothenomys eleusis | Eothenomys | Cricetidae | China, Asia | tissue (liver) |
| S2914 | Eothenomys eleusis | Eothenomys | Cricetidae | China, Asia | tissue (liver) |
| S2905 | Eothenomys eleusis | Eothenomys | Cricetidae | China, Asia | tissue (liver) |
| S2657 | Eothenomys melanogaster | Eothenomys | Cricetidae | China, Asia | tissue (liver) |
| S2763 | Eothenomys melanogaster | Eothenomys | Cricetidae | China, Asia | tissue (liver) |
| S2645 | Eothenomys melanogaster | Eothenomys | Cricetidae | China, Asia | tissue (liver) |
| S2619 | Eothenomys melanogaster | Eothenomys | Cricetidae | China, Asia | tissue (liver) |
| S2959 | Eothenomys miletus | Eothenomys | Cricetidae | China, Asia | tissue (liver) |
| S2970 | Eothenomys miletus | Eothenomys | Cricetidae | China, Asia | tissue (liver) |
| COVROD 3567 | Acomys cahirinus | Acomys | Muridae | Saudi Arabia, Asia | RNA (liver) |
| COVROD 3564 | Acomys dimidiatus | Acomys | Muridae | Saudi Arabia, Asia | RNA (liver) |
| COVROD 3570 | Acomys dimidiatus | Acomys | Muridae | Saudi Arabia, Asia | RNA (liver) |
| COVROD 3590 | Acomys dimidiatus | Acomys | Muridae | Saudi Arabia, Asia | RNA (liver) |
| COVROD 3562 | Acomys dimidiatus | Acomys | Muridae | Saudi Arabia, Asia | RNA (liver) |
| COVROD 3559 | Dipodillus dasyurus | Dipodillus | Muridae | Saudi Arabia, Asia | RNA (liver) |
| COVROD 3589 | Dipodillus dasyurus | Dipodillus | Muridae | Saudi Arabia, Asia | RNA (liver) |
| COVROD 3580 | Dipodillus dasyurus | Dipodillus | Muridae | Saudi Arabia, Asia | RNA (liver) |
| COVROD 3581 | Dipodillus dasyurus | Dipodillus | Muridae | Saudi Arabia, Asia | RNA (liver) |
| PL52-UNIKIS48 | Hylomyscus sp. | Hylomyscus | Muridae | DRC, Africa | DBS |
| PL53-UNIKIS49 | Hylomyscus sp. | Hylomyscus | Muridae | DRC, Africa | DBS |
| TA289 | Lophuromys laticeps | Lophuromys | Muridae | Tanzania, Africa | tissue (liver) |
| AFGN30 | Lophuromys sikapusi | Lophuromys | Muridae | Guinea, Africa | tissue (liver) |
| AFGN46 | Lophuromys sikapusi | Lophuromys | Muridae | Guinea, Africa | tissue (liver) |
| AFGN47 | Lophuromys sikapusi | Lophuromys | Muridae | Guinea, Africa | tissue (liver) |
| TA109 | Lophuromys stanleyi | Lophuromys | Muridae | Tanzania, Africa | tissue (liver) |
| PL65-UNIKIS75 | Lophyromys sp. | Lophuromys | Muridae | DRC, Africa | DBS |
| PL120-UNIKIS134 | Lophyromys huttereri | Lophuromys | Muridae | DRC, Africa | DBS |
| ABA101 | Meriones libycus | Meriones | Muridae | Saudi Arabia, Asia | RNA (plasma) |
| ABA117 | Meriones libycus | Meriones | Muridae | Saudi Arabia, Asia | RNA (plasma) |
| ABA123 | Meriones libycus | Meriones | Muridae | Saudi Arabia, Asia | RNA (plasma) |
| ABA32 | Meriones libycus | Meriones | Muridae | Saudi Arabia, Asia | RNA (plasma) |
| ABA55 | Meriones libycus | Meriones | Muridae | Saudi Arabia, Asia | RNA (plasma) |
| ABA62 | Meriones libycus | Meriones | Muridae | Saudi Arabia, Asia | RNA (plasma) |
| ABA70 | Meriones libycus | Meriones | Muridae | Saudi Arabia, Asia | RNA (plasma) |
| ABA5 | Meriones libycus | Meriones | Muridae | Saudi Arabia, Asia | RNA (plasma) |
| ABA76 | Meriones libycus | Meriones | Muridae | Saudi Arabia, Asia | RNA (plasma) |
| ABA81 | Meriones libycus | Meriones | Muridae | Saudi Arabia, Asia | RNA (plasma) |
| ABA99 | Meriones libycus | Meriones | Muridae | Saudi Arabia, Asia | RNA (plasma) |
| ABA13 | Meriones libycus | Meriones | Muridae | Saudi Arabia, Asia | RNA (plasma) |
| ABA15 | Meriones libycus | Meriones | Muridae | Saudi Arabia, Asia | RNA (plasma) |
| ABA18 | Meriones libycus | Meriones | Muridae | Saudi Arabia, Asia | RNA (plasma) |
| ABA97 | Meriones libycus | Meriones | Muridae | Saudi Arabia, Asia | RNA (plasma) |
| COVROD 3561 | Meriones rex | Meriones | Muridae | Saudi Arabia, Asia | RNA (liver) |
| COVROD 3598 | Meriones rex | Meriones | Muridae | Saudi Arabia, Asia | RNA (liver) |
| COVROD 3563 | Meriones rex | Meriones | Muridae | Saudi Arabia, Asia | RNA (liver) |
| S2741 | Micromys minutus | Micromys | Muridae | China, Asia | tissue (liver) |
| S2655 | Niviventer confucianus | Niviventer | Muridae | China, Asia | tissue (liver) |
| TA156 | Praomys jacksoni | Praomys | Muridae | Tanzania, Africa | tissue (liver) |
| S2889 | Rattus andamanensis | Rattus | Muridae | China, Asia | tissue (liver) |
| COVROD 3565 | Rattus rattus | Rattus | Muridae | Saudi Arabia, Asia | RNA (liver) |
| **Hepacivirus positive sample been detected and sequenced by our collaborators** | | | | | |
| CZ21 | Clethrionomys glareolus | Clethrionomys | Cricetidae | Czech Republic, Europe | RNAseq raw data |
